## Supplementary figures and images for "Distinct transcriptomic profile of satellite cells contributes to preservation of neuromuscular junctions in extraocular muscles of ALS mice"

### Figure 1-figure supplement 1

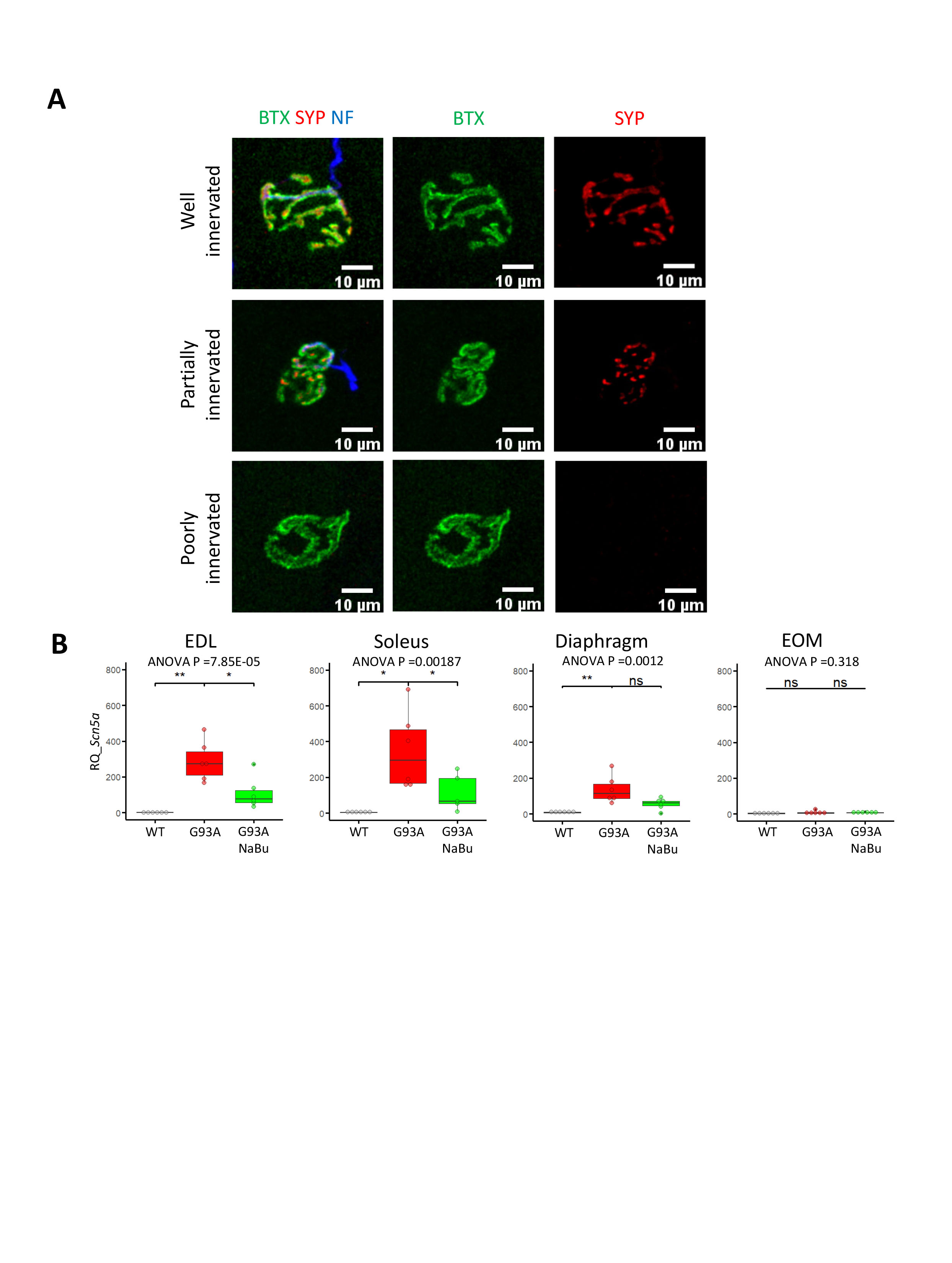

### Figure 1-figure supplement 2

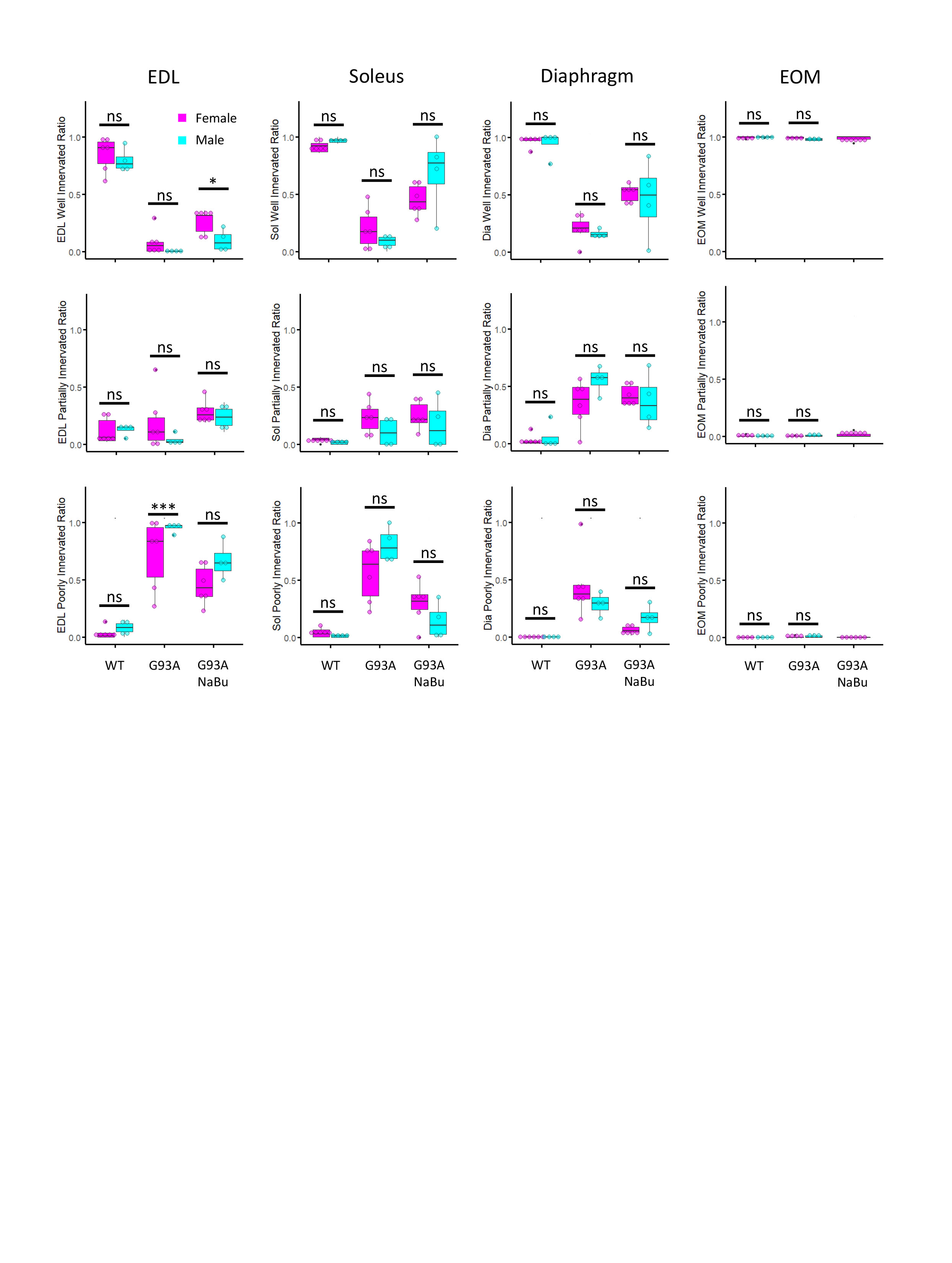

### Figure 3-figure supplement 1

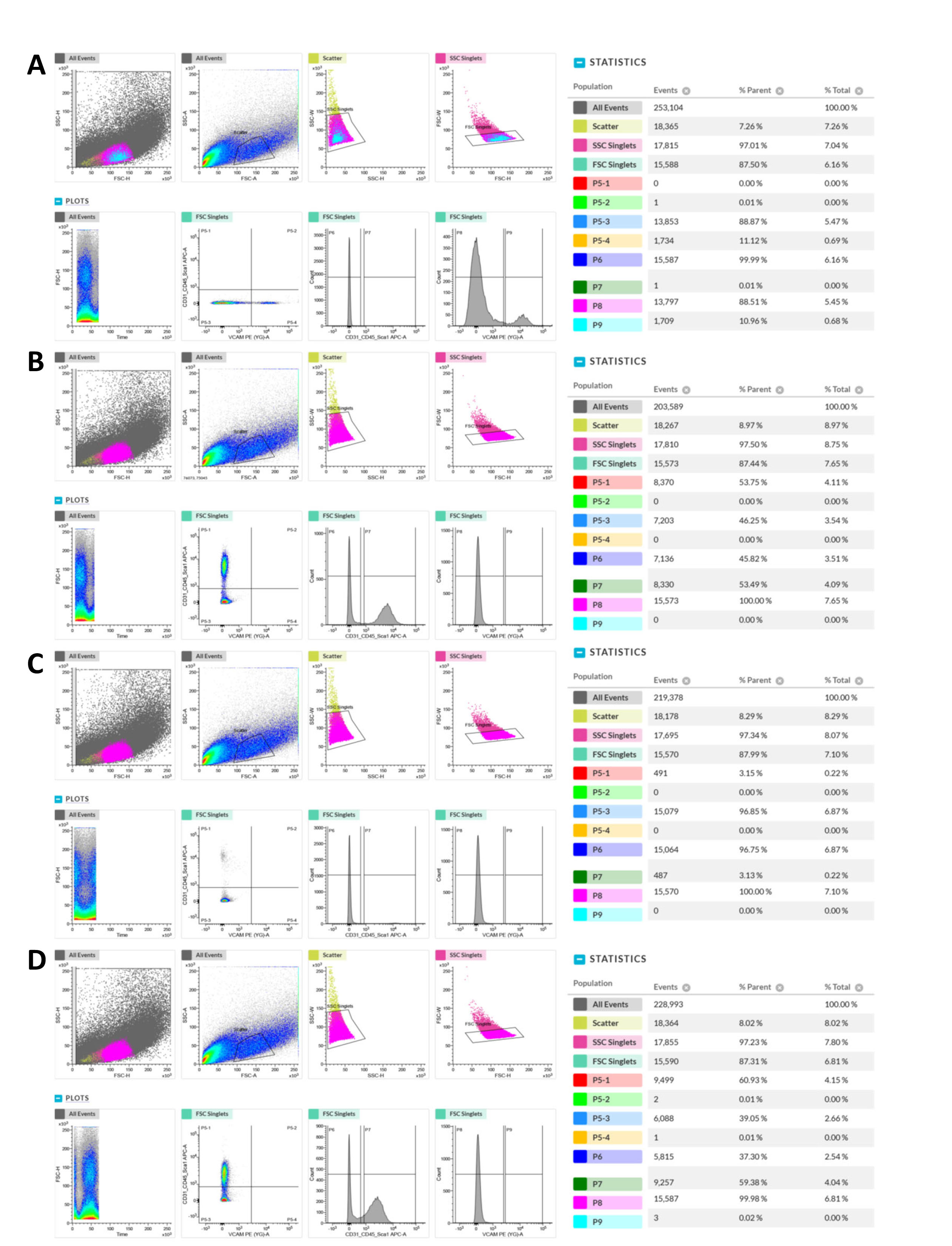

### Figure 3-figure supplement 2

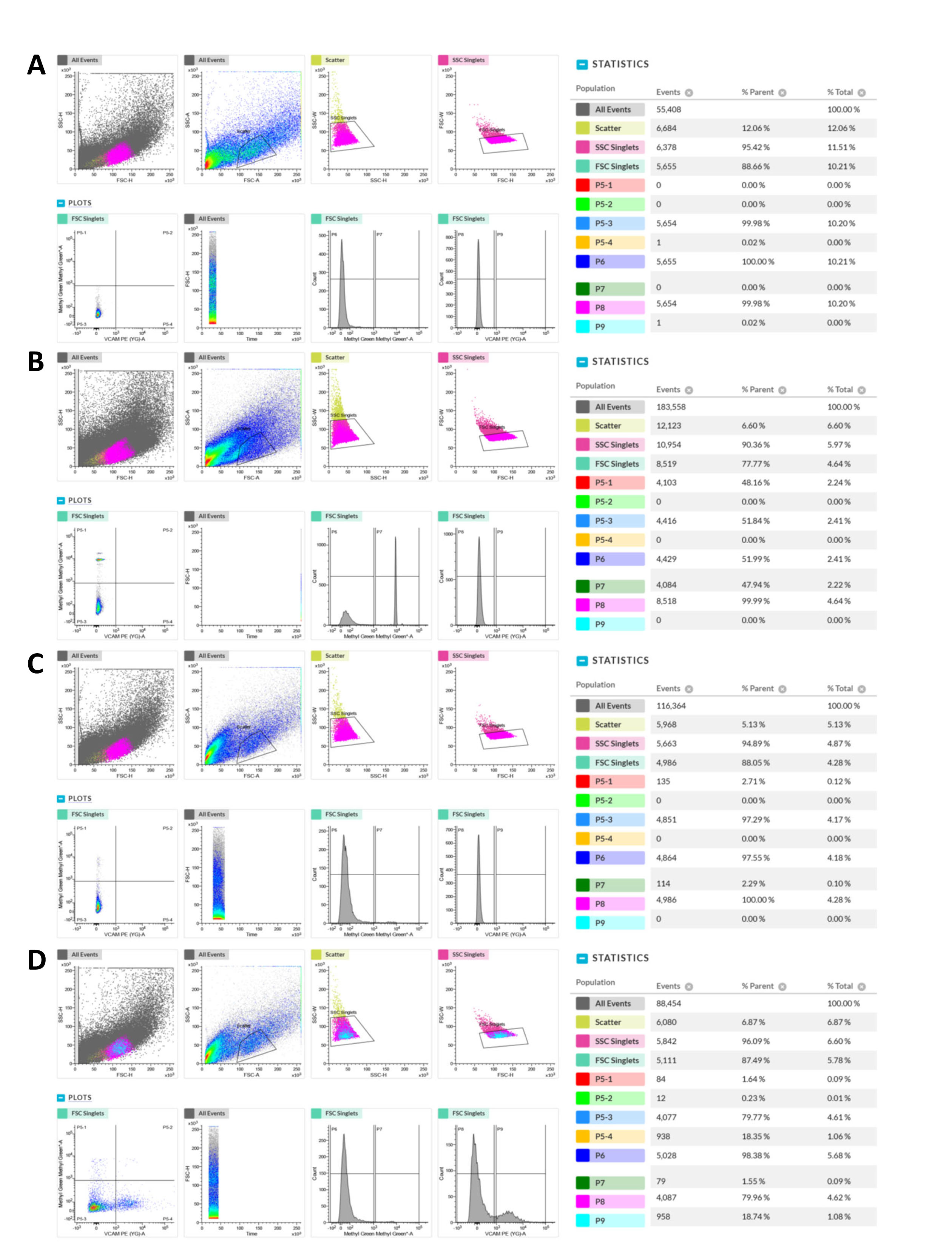

### Figure 4-figure supplement 1

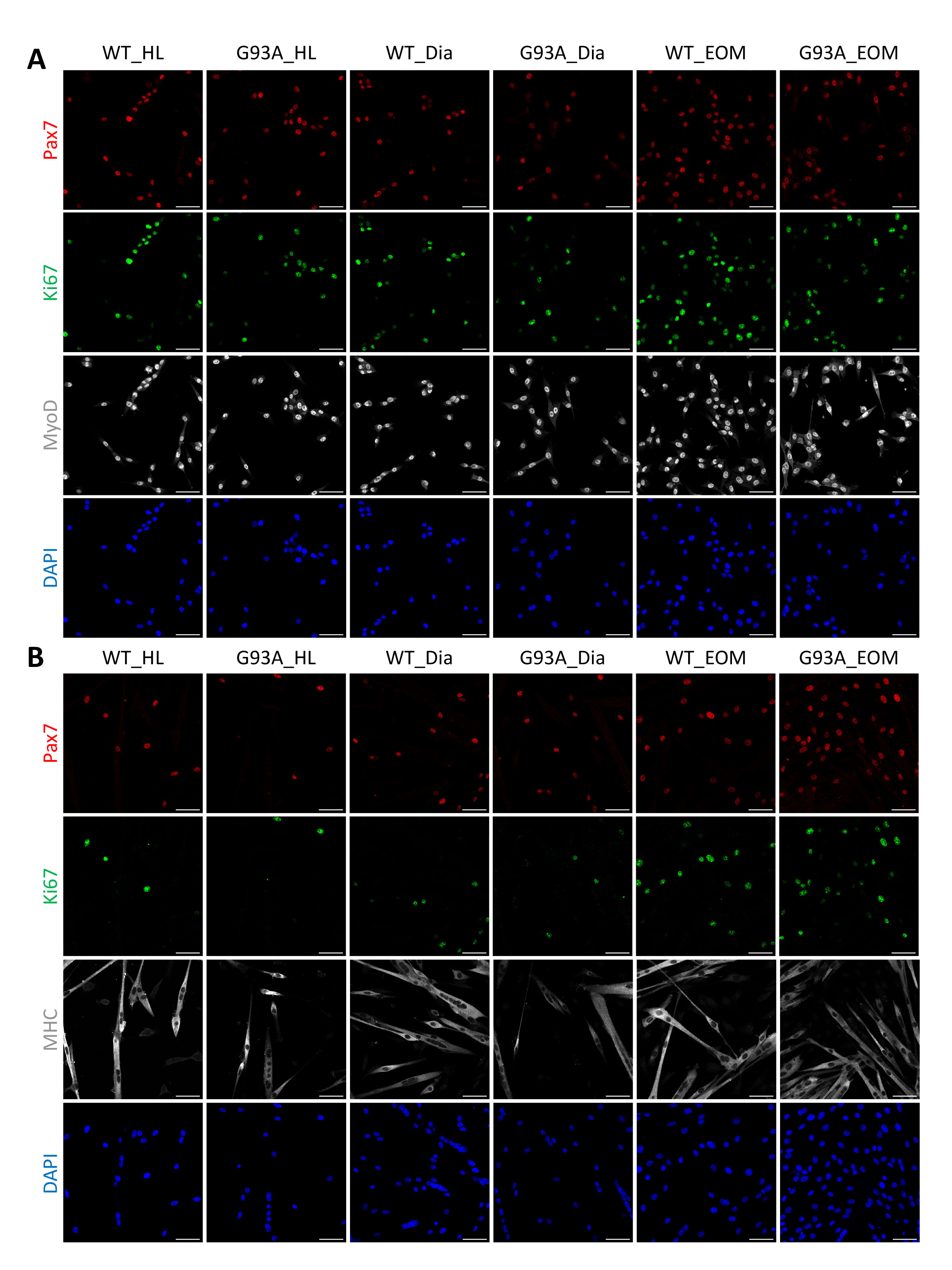

### Figure 4-figure supplement 2

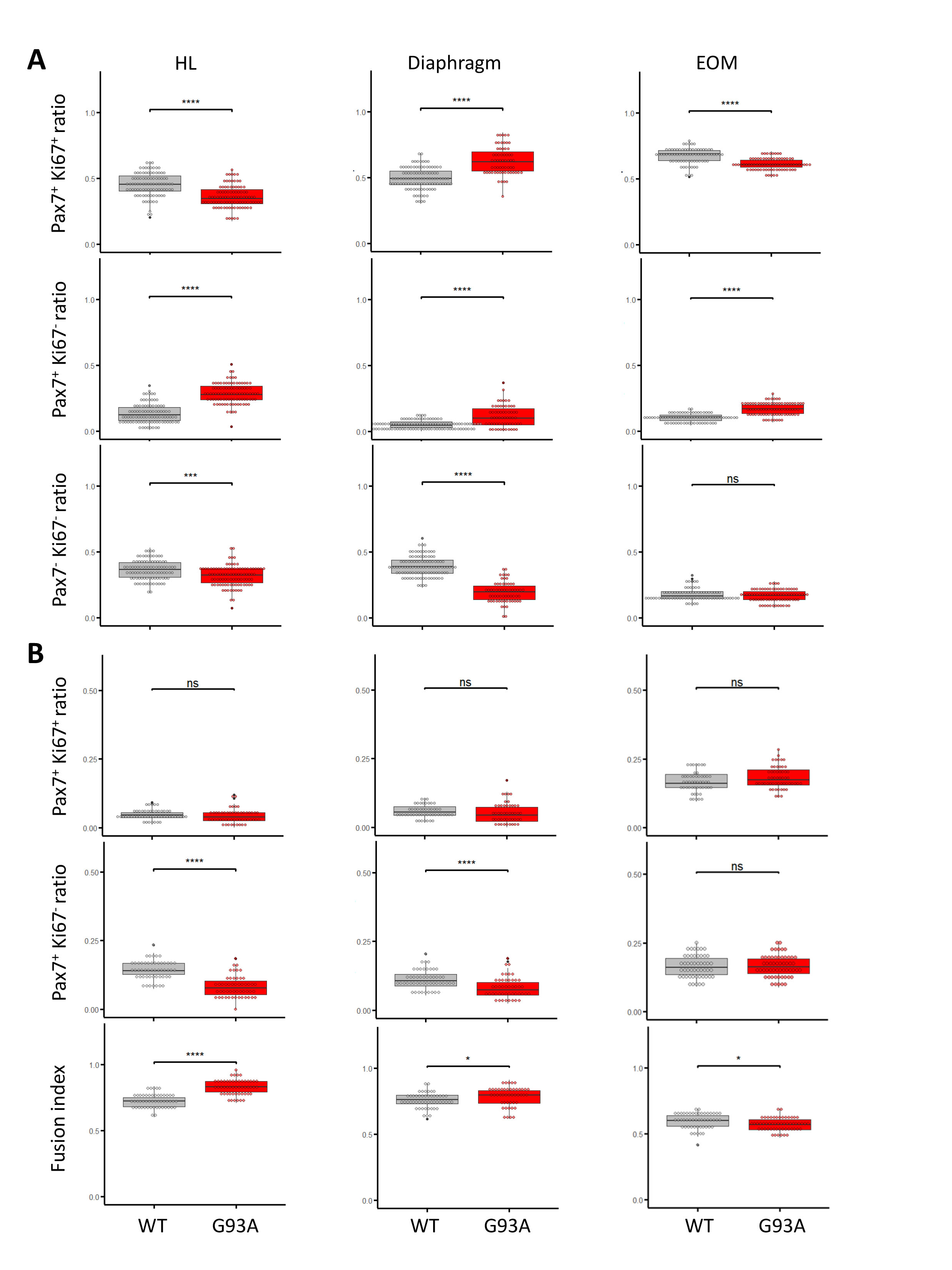

### Figure 6-figure supplement 1

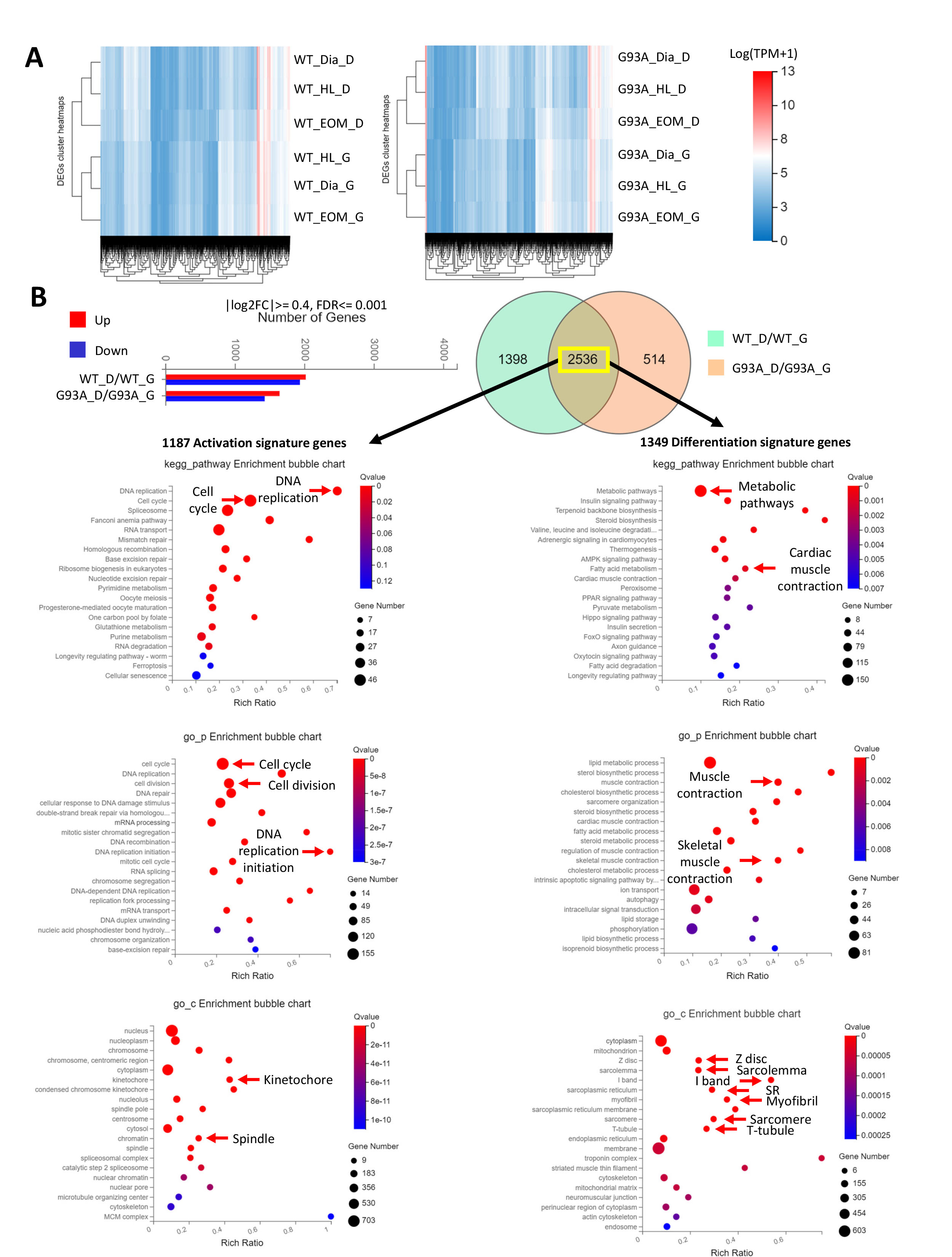

### Figure 7-figure supplement 1

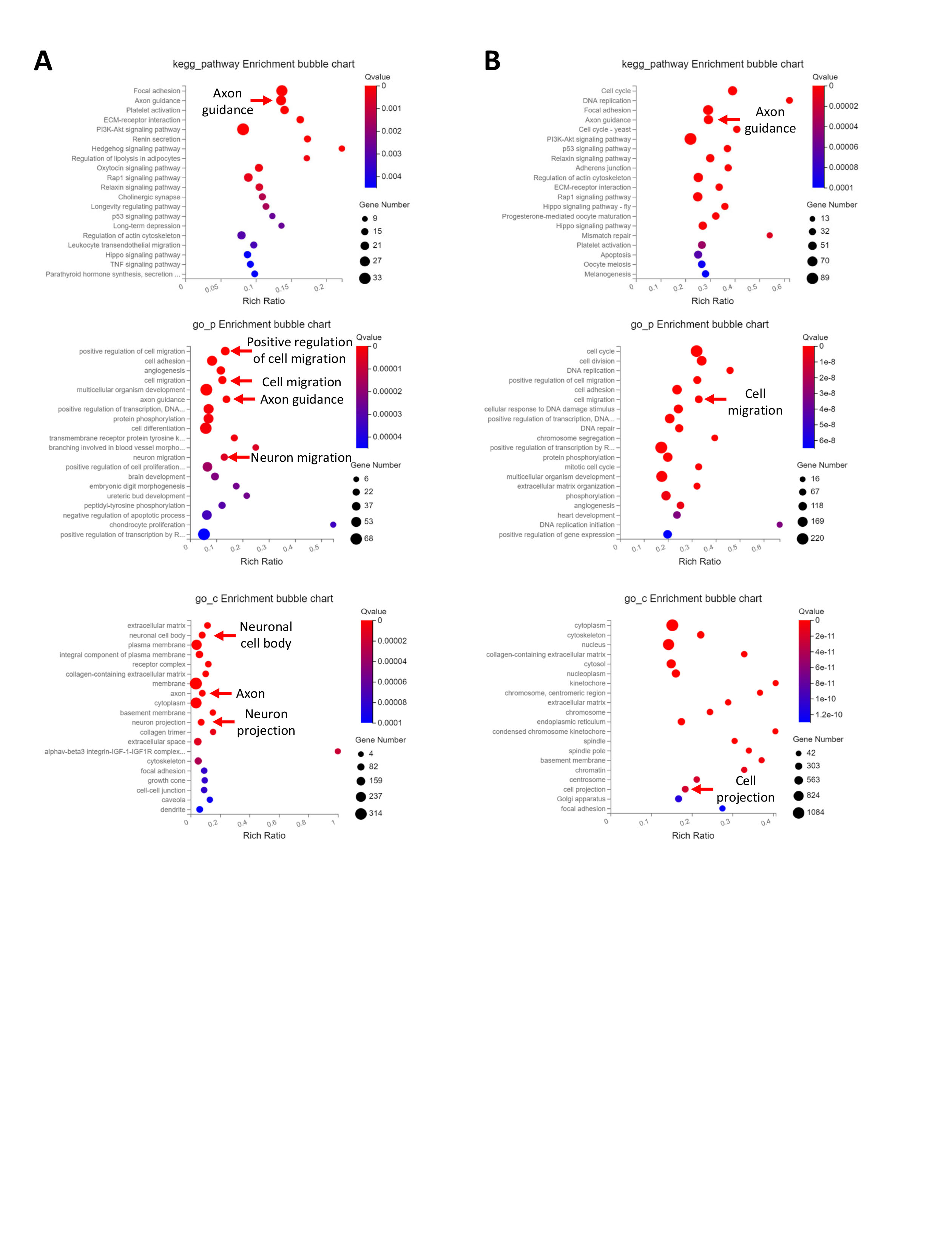

### Figure 8-figure supplement 1

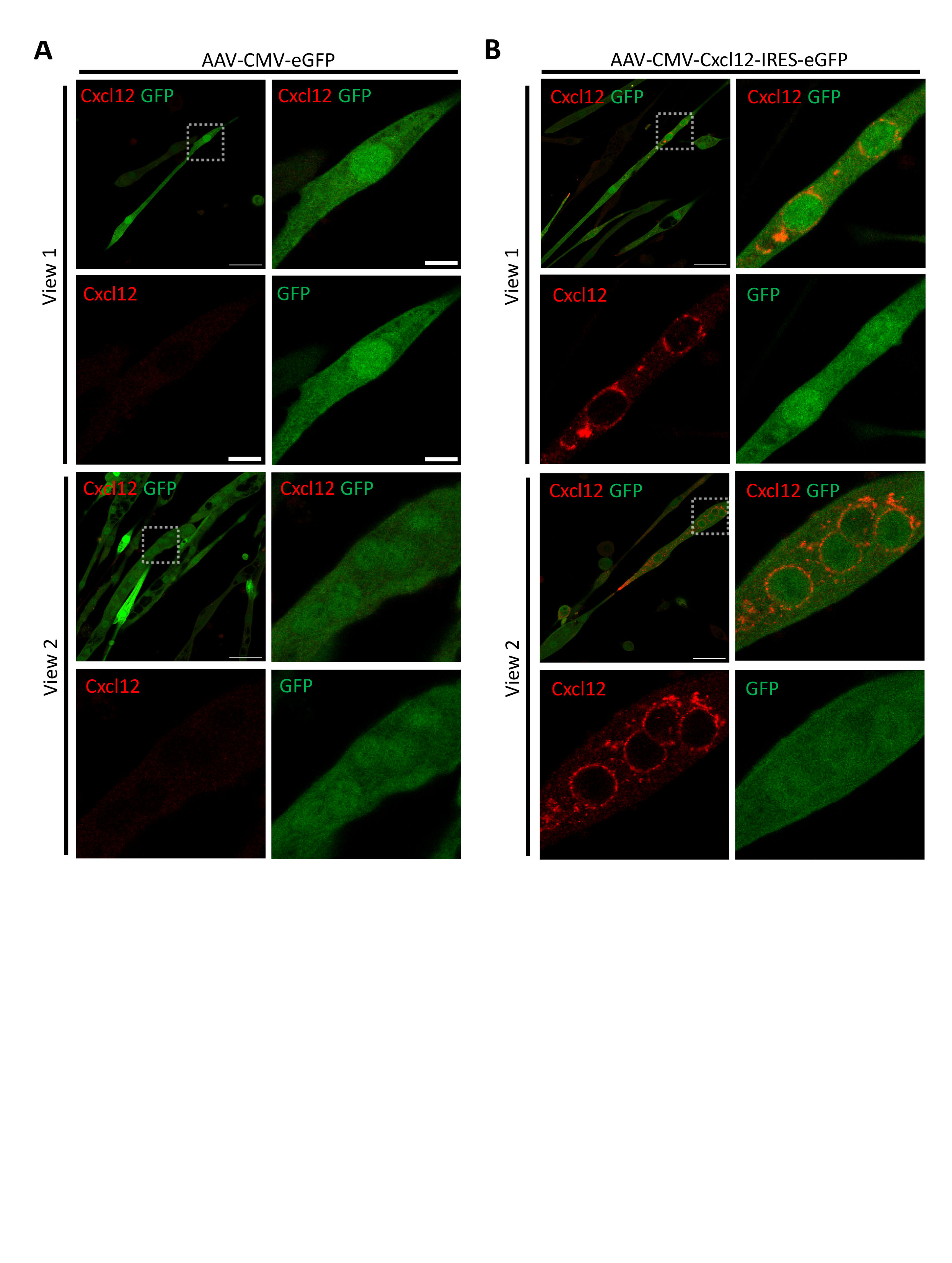

### Figure 8-figure supplement 2

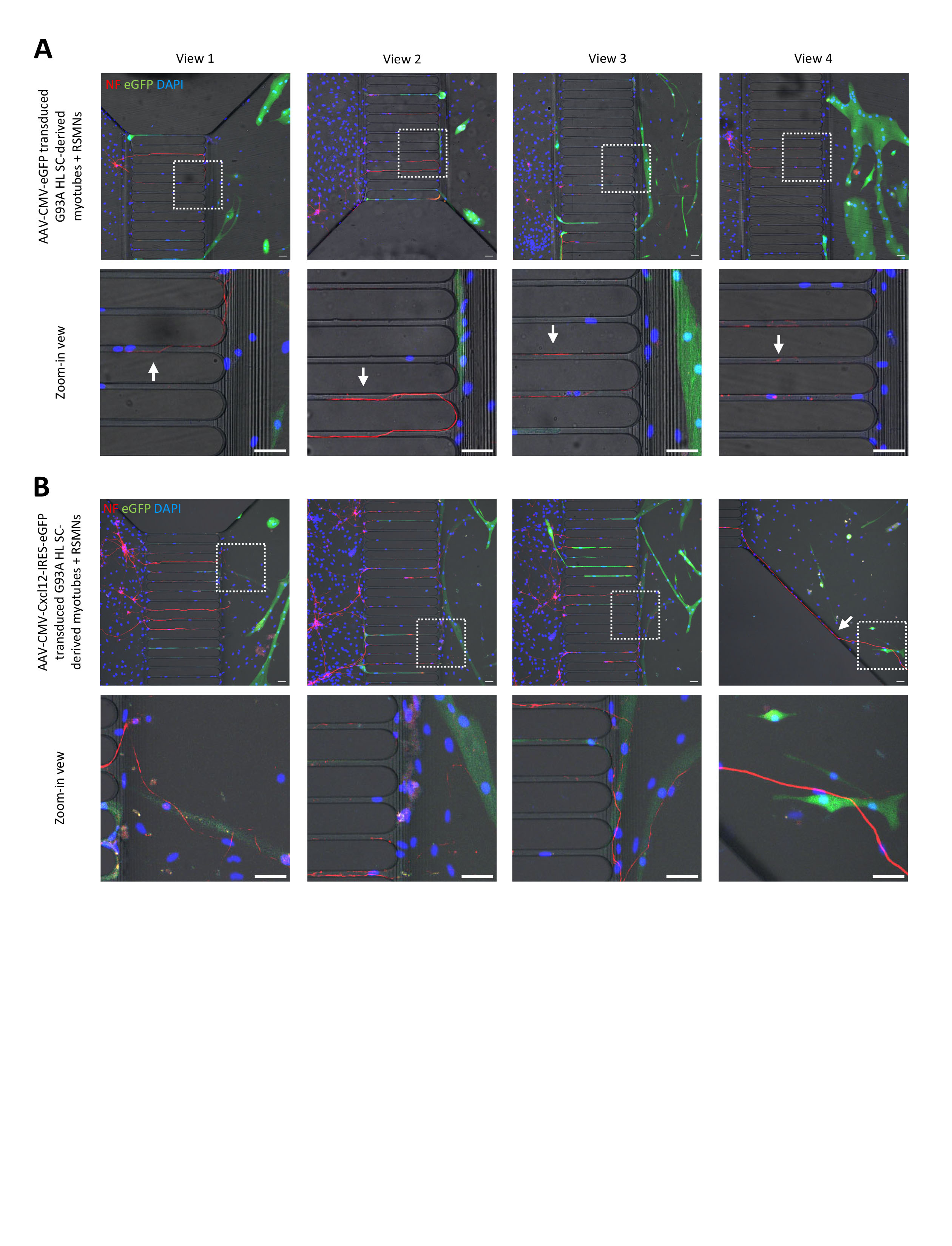

### Figure 9-figure supplement 1

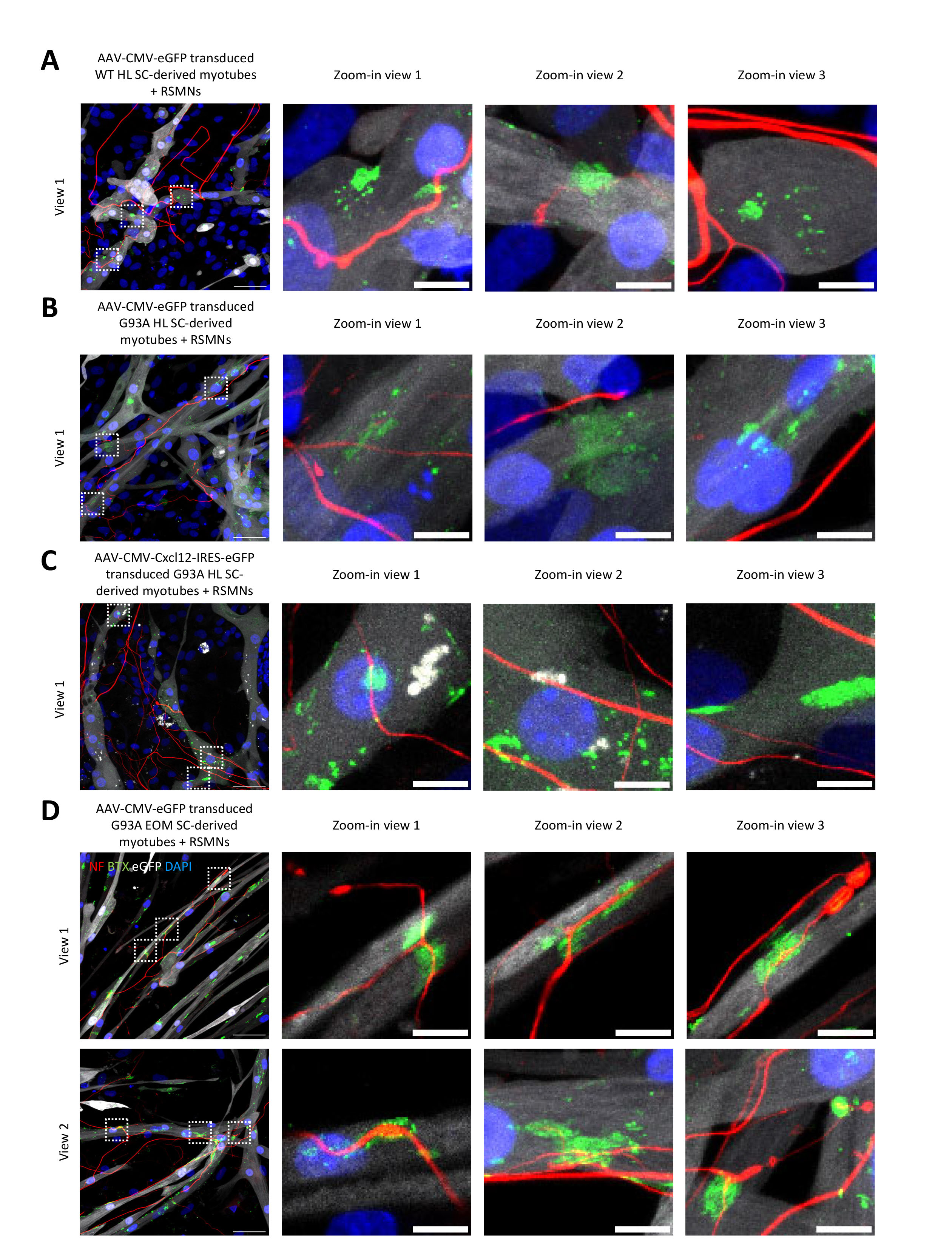
