## Supplementary material for "Distinct transcriptomic profile of satellite cells contributes to preservation of neuromuscular junctions in extraocular muscles of ALS mice": Figure 1-Source Data 1

**Figure 1-Source Data 1. Quantification results of the three types of NMJs in muscles of different origins and treatment conditions.**

| Mouse | Gender | Muscle | Group | Well | Partial | Poor | Well  ratio | Partial  ratio | Poor  ratio |
| --- | --- | --- | --- | --- | --- | --- | --- | --- | --- |
| EDL-1 | M | EDL | WT | 42 | 9 | 9 | 0.7 | 0.15 | 0.15 |
| EDL-2 | F | EDL | WT | 72 | 3 | 3 | 0.923077 | 0.038462 | 0.038462 |
| EDL-3 | F | EDL | WT | 52 | 1 | 0 | 0.981132 | 0.018868 | 0 |
| EDL-4 | F | EDL | WT | 32 | 3 | 1 | 0.888889 | 0.083333 | 0.027778 |
| EDL-5 | M | EDL | WT | 64 | 12 | 5 | 0.790123 | 0.148148 | 0.061728 |
| EDL-6 | M | EDL | WT | 18 | 1 | 0 | 0.947368 | 0.052632 | 0 |
| EDL-7 | M | EDL | WT | 48 | 10 | 7 | 0.738462 | 0.153846 | 0.107692 |
| EDL-8 | F | EDL | WT | 53 | 20 | 0 | 0.726027 | 0.273973 | 0 |
| EDL-9 | F | EDL | WT | 32 | 1 | 0 | 0.969697 | 0.030303 | 0 |
| EDL-10 | F | EDL | WT | 91 | 37 | 20 | 0.614865 | 0.25 | 0.135135 |
| EDL-11 | F | EDL | G93A | 3 | 14 | 110 | 0.023622 | 0.110236 | 0.866142 |
| EDL-12 | F | EDL | G93A | 3 | 24 | 10 | 0.081081 | 0.648649 | 0.27027 |
| EDL-13 | F | EDL | G93A | 0 | 0 | 28 | 0 | 0 | 1 |
| EDL-14 | M | EDL | G93A | 0 | 1 | 40 | 0 | 0.02439 | 0.97561 |
| EDL-15 | M | EDL | G93A | 0 | 6 | 49 | 0 | 0.109091 | 0.890909 |
| EDL-16 | M | EDL | G93A | 1 | 3 | 152 | 0.00641 | 0.019231 | 0.974359 |
| EDL-17 | M | EDL | G93A | 1 | 1 | 80 | 0.012195 | 0.012195 | 0.97561 |
| EDL-18 | F | EDL | G93A | 17 | 16 | 25 | 0.293103 | 0.275862 | 0.431034 |
| EDL-19 | F | EDL | G93A | 8 | 10 | 75 | 0.086022 | 0.107527 | 0.806452 |
| EDL-20 | F | EDL | G93A | 0 | 1 | 80 | 0 | 0.012346 | 0.987654 |
| EDL-21 | F | EDL | G93A_NaBu | 39 | 30 | 40 | 0.357798 | 0.275229 | 0.366972 |
| EDL-22 | M | EDL | G93A_NaBu | 2 | 30 | 50 | 0.02439 | 0.365854 | 0.609756 |
| EDL-23 | M | EDL | G93A_NaBu | 22 | 29 | 50 | 0.217822 | 0.287129 | 0.49505 |
| EDL-24 | F | EDL | G93A_NaBu | 18 | 26 | 13 | 0.315789 | 0.45614 | 0.22807 |
| EDL-25 | F | EDL | G93A_NaBu | 18 | 19 | 20 | 0.315789 | 0.333333 | 0.350877 |
| EDL-26 | M | EDL | G93A_NaBu | 14 | 20 | 74 | 0.12963 | 0.185185 | 0.685185 |
| EDL-27 | M | EDL | G93A_NaBu | 1 | 6 | 49 | 0.017857 | 0.107143 | 0.875 |
| EDL-28 | F | EDL | G93A_NaBu | 22 | 13 | 34 | 0.318841 | 0.188406 | 0.492754 |
| EDL-29 | F | EDL | G93A_NaBu | 15 | 27 | 72 | 0.131579 | 0.236842 | 0.631579 |
| EDL-30 | F | EDL | G93A_NaBu | 8 | 13 | 43 | 0.125 | 0.203125 | 0.671875 |
| Sol-1 | M | Sol | WT | 42 | 0 | 1 | 0.976744 | 0 | 0.023256 |
| Sol-2 | F | Sol | WT | 47 | 3 | 2 | 0.903846 | 0.057692 | 0.038462 |
| Sol-3 | M | Sol | WT | 44 | 0 | 0 | 1 | 0 | 0 |
| Sol-4 | M | Sol | WT | 48 | 2 | 0 | 0.96 | 0.04 | 0 |
| Sol-5 | M | Sol | WT | 32 | 1 | 1 | 0.941176 | 0.029412 | 0.029412 |
| Sol-6 | F | Sol | WT | 18 | 1 | 0 | 0.947368 | 0.052632 | 0 |
| Sol-7 | F | Sol | WT | 55 | 4 | 5 | 0.859375 | 0.0625 | 0.078125 |
| Sol-8 | F | Sol | WT | 74 | 3 | 2 | 0.936709 | 0.037975 | 0.025316 |
| Sol-9 | F | Sol | WT | 29 | 0 | 0 | 1 | 0 | 0 |
| Sol-10 | F | Sol | WT | 25 | 1 | 3 | 0.862069 | 0.034483 | 0.103448 |
| Sol-11 | F | Sol | G93A | 6 | 13 | 21 | 0.15 | 0.325 | 0.525 |
| Sol-12 | F | Sol | G93A | 14 | 18 | 9 | 0.341463 | 0.439024 | 0.219512 |
| Sol-13 | F | Sol | G93A | 20 | 9 | 13 | 0.47619 | 0.214286 | 0.309524 |
| Sol-14 | M | Sol | G93A | 0 | 0 | 15 | 0 | 0 | 1 |
| Sol-15 | F | Sol | G93A | 0 | 2 | 6 | 0 | 0.25 | 0.75 |
| Sol-16 | M | Sol | G93A | 4 | 12 | 36 | 0.076923 | 0.230769 | 0.692308 |
| Sol-17 | M | Sol | G93A | 2 | 0 | 13 | 0.133333 | 0 | 0.866667 |
| Sol-18 | M | Sol | G93A | 5 | 8 | 27 | 0.125 | 0.2 | 0.675 |
| Sol-19 | F | Sol | G93A | 2 | 5 | 37 | 0.045455 | 0.113636 | 0.840909 |
| Sol-20 | F | Sol | G93A | 9 | 2 | 35 | 0.195652 | 0.043478 | 0.76087 |
| Sol-21 | F | Sol | G93A_NaBu | 46 | 19 | 30 | 0.484211 | 0.2 | 0.315789 |
| Sol-22 | F | Sol | G93A_NaBu | 14 | 9 | 15 | 0.368421 | 0.236842 | 0.394737 |
| Sol-23 | M | Sol | G93A_NaBu | 18 | 6 | 1 | 0.72 | 0.24 | 0.04 |
| Sol-24 | M | Sol | G93A_NaBu | 8 | 18 | 14 | 0.2 | 0.45 | 0.35 |
| Sol-25 | M | Sol | G93A_NaBu | 28 | 0 | 6 | 0.823529 | 0 | 0.176471 |
| Sol-26 | M | Sol | G93A_NaBu | 5 | 0 | 0 | 1 | 0 | 0 |
| Sol-27 | F | Sol | G93A_NaBu | 13 | 3 | 18 | 0.382353 | 0.088235 | 0.529412 |
| Sol-28 | F | Sol | G93A_NaBu | 19 | 6 | 7 | 0.59375 | 0.1875 | 0.21875 |
| Sol-29 | F | Sol | G93A_NaBu | 13 | 19 | 15 | 0.276596 | 0.404255 | 0.319149 |
| Sol-30 | F | Sol | G93A_NaBu | 8 | 5 | 0 | 0.615385 | 0.384615 | 0 |
| Dia-1 | M | Dia | WT | 63 | 19 | 0 | 0.768293 | 0.231707 | 0 |
| Dia-2 | F | Dia | WT | 118 | 17 | 0 | 0.874074 | 0.125926 | 0 |
| Dia-3 | F | Dia | WT | 176 | 1 | 0 | 0.99435 | 0.00565 | 0 |
| Dia-4 | F | Dia | WT | 87 | 1 | 0 | 0.988636 | 0.011364 | 0 |
| Dia-5 | M | Dia | WT | 177 | 0 | 0 | 1 | 0 | 0 |
| Dia-6 | M | Dia | WT | 222 | 0 | 0 | 1 | 0 | 0 |
| Dia-7 | M | Dia | WT | 105 | 0 | 0 | 1 | 0 | 0 |
| Dia-8 | F | Dia | WT | 146 | 0 | 0 | 1 | 0 | 0 |
| Dia-9 | F | Dia | WT | 124 | 4 | 0 | 0.96875 | 0.03125 | 0 |
| Dia-10 | F | Dia | WT | 111 | 1 | 0 | 0.991071 | 0.008929 | 0 |
| Dia-11 | F | Dia | G93A | 0 | 2 | 139 | 0 | 0.014184 | 0.985816 |
| Dia-12 | F | Dia | G93A | 42 | 84 | 23 | 0.281879 | 0.563758 | 0.154362 |
| Dia-13 | F | Dia | G93A | 16 | 50 | 32 | 0.163265 | 0.510204 | 0.326531 |
| Dia-14 | M | Dia | G93A | 25 | 108 | 47 | 0.138889 | 0.6 | 0.261111 |
| Dia-15 | M | Dia | G93A | 15 | 62 | 15 | 0.163043 | 0.673913 | 0.163043 |
| Dia-16 | F | Dia | G93A | 63 | 41 | 72 | 0.357955 | 0.232955 | 0.409091 |
| Dia-17 | M | Dia | G93A | 18 | 83 | 50 | 0.119205 | 0.549669 | 0.331126 |
| Dia-18 | M | Dia | G93A | 25 | 47 | 47 | 0.210084 | 0.394958 | 0.394958 |
| Dia-19 | F | Dia | G93A | 9 | 15 | 21 | 0.2 | 0.333333 | 0.466667 |
| Dia-20 | F | Dia | G93A | 20 | 41 | 32 | 0.215054 | 0.44086 | 0.344086 |
| Dia-21 | F | Dia | G93A_NaBu | 78 | 47 | 13 | 0.565217 | 0.34058 | 0.094203 |
| Dia-22 | F | Dia | G93A_NaBu | 86 | 61 | 17 | 0.52439 | 0.371951 | 0.103659 |
| Dia-23 | F | Dia | G93A_NaBu | 97 | 74 | 2 | 0.560694 | 0.427746 | 0.011561 |
| Dia-24 | M | Dia | G93A_NaBu | 1 | 49 | 22 | 0.013889 | 0.680556 | 0.305556 |
| Dia-25 | M | Dia | G93A_NaBu | 48 | 51 | 19 | 0.40678 | 0.432203 | 0.161017 |
| Dia-26 | M | Dia | G93A_NaBu | 45 | 18 | 14 | 0.584416 | 0.233766 | 0.181818 |
| Dia-27 | M | Dia | G93A_NaBu | 61 | 10 | 2 | 0.835616 | 0.136986 | 0.027397 |
| Dia-28 | F | Dia | G93A_NaBu | 90 | 49 | 9 | 0.608108 | 0.331081 | 0.060811 |
| Dia-29 | F | Dia | G93A_NaBu | 79 | 99 | 8 | 0.424731 | 0.532258 | 0.043011 |
| Dia-30 | F | Dia | G93A_NaBu | 25 | 31 | 3 | 0.423729 | 0.525424 | 0.050847 |
| EOM-1 | M | EOM | WT | 82 | 0 | 0 | 1 | 0 | 0 |
| EOM-2 | F | EOM | WT | 39 | 0 | 0 | 1 | 0 | 0 |
| EOM-3 | M | EOM | WT | 33 | 0 | 0 | 1 | 0 | 0 |
| EOM-4 | M | EOM | WT | 157 | 1 | 0 | 0.993671 | 0.006329 | 0 |
| EOM-5 | M | EOM | WT | 62 | 0 | 0 | 1 | 0 | 0 |
| EOM-6 | F | EOM | WT | 94 | 2 | 0 | 0.979167 | 0.020833 | 0 |
| EOM-7 | F | EOM | WT | 87 | 0 | 0 | 1 | 0 | 0 |
| EOM-8 | F | EOM | WT | 63 | 0 | 0 | 1 | 0 | 0 |
| EOM-9 | M | EOM | G93A | 86 | 2 | 0 | 0.977273 | 0.022727 | 0 |
| EOM-10 | M | EOM | G93A | 160 | 0 | 6 | 0.963855 | 0 | 0.036145 |
| EOM-11 | F | EOM | G93A | 88 | 0 | 2 | 0.977778 | 0 | 0.022222 |
| EOM-12 | M | EOM | G93A | 74 | 0 | 0 | 1 | 0 | 0 |
| EOM-13 | F | EOM | G93A | 120 | 1 | 0 | 0.991736 | 0.008264 | 0 |
| EOM-14 | F | EOM | G93A | 66 | 0 | 0 | 1 | 0 | 0 |
| EOM-15 | F | EOM | G93A | 90 | 0 | 0 | 1 | 0 | 0 |
| EOM-16 | F | EOM | G93A_NaBu | 102 | 0 | 0 | 1 | 0 | 0 |
| EOM-17 | F | EOM | G93A_NaBu | 143 | 0 | 0 | 1 | 0 | 0 |
| EOM-18 | F | EOM | G93A_NaBu | 232 | 0 | 0 | 1 | 0 | 0 |
| EOM-19 | F | EOM | G93A_NaBu | 37 | 0 | 0 | 1 | 0 | 0 |
| EOM-20 | F | EOM | G93A_NaBu | 36 | 1 | 0 | 0.972973 | 0.027027 | 0 |
| EOM-21 | F | EOM | G93A_NaBu | 34 | 2 | 0 | 0.944444 | 0.055556 | 0 |
