## Supplementary material for "Distinct transcriptomic profile of satellite cells contributes to preservation of neuromuscular junctions in extraocular muscles of ALS mice": Figure 1-Source Data 2

**Figure 1-Source Data 2. qRT-PCR results for *Scn5a* relative expression in whole muscles of different origins and treatment conditions.** The calculation of ddCt used the averaged dCt value of *Scn5a* of EDL muscles derived from WT mice as the normalization control.

| Mouse | Gender | Muscle | Group | Ct_*Scn5a* | Ct_*Gapdh* | dCT_*Scn5a* | ddCT_*Scn5a* | RQ_*Scn5a* |
| --- | --- | --- | --- | --- | --- | --- | --- | --- |
| EDL-1 | F | EDL | WT | 33.74081 | 17.48398 | 16.25683 | -0.33644 | 1.262639 |
| EDL-2 | F | EDL | WT | 35.00173 | 17.88853 | 17.1132 | 0.519928 | 0.697407 |
| EDL-3 | M | EDL | WT | 34.14015 | 16.88818 | 17.25197 | 0.658702 | 0.633448 |
| EDL-4 | M | EDL | WT | 33.68679 | 16.8915 | 16.79529 | 0.202021 | 0.869332 |
| EDL-5 | F | EDL | WT | 33.26038 | 17.31091 | 15.94947 | -0.6438 | 1.562442 |
| EDL-6 | M | EDL | WT | 32.76152 | 16.56865 | 16.19287 | -0.4004 | 1.319869 |
| EDL-7 | F | EDL | G93A | 25.97019 | 17.48626 | 8.483927 | -8.10934 | 276.1566 |
| EDL-8 | F | EDL | G93A | 25.40596 | 17.67061 | 7.73535 | -8.85792 | 463.9805 |
| EDL-9 | M | EDL | G93A | 25.94594 | 16.73245 | 9.213488 | -7.37978 | 166.5466 |
| EDL-10 | M | EDL | G93A | 25.81356 | 16.79055 | 9.02301 | -7.57026 | 190.0533 |
| EDL-11 | F | EDL | G93A | 24.90046 | 16.81133 | 8.089125 | -8.50415 | 363.0804 |
| EDL-12 | M | EDL | G93A | 24.35285 | 15.82312 | 8.529735 | -8.06354 | 267.526 |
| EDL-13 | F | EDL | G93A_NaBu | 27.95392 | 17.82598 | 10.12794 | -6.46533 | 88.36065 |
| EDL-14 | M | EDL | G93A_NaBu | 28.25739 | 17.71075 | 10.54665 | -6.04662 | 66.10209 |
| EDL-15 | F | EDL | G93A_NaBu | 28.14194 | 18.65344 | 9.4885 | -7.10477 | 137.6413 |
| EDL-16 | F | EDL | G93A_NaBu | 30.95402 | 20.10743 | 10.84659 | -5.74668 | 53.69352 |
| EDL-17 | M | EDL | G93A_NaBu | 24.42928 | 15.91946 | 8.50982 | -8.08345 | 271.2445 |
| EDL-18 | M | EDL | G93A_NaBu | 21.0703 | 16.1138 | 11.55161 | -5.04166 | 32.9375 |
| Sol-1 | F | Sol | WT | 33.37701 | 18.71398 | 14.66302 | -1.93025 | 3.811201 |
| Sol-2 | F | Sol | WT | 32.85528 | 18.12337 | 14.7319 | -1.86137 | 3.633521 |
| Sol-3 | M | Sol | WT | 34.11594 | 18.94196 | 15.17398 | -1.41929 | 2.674538 |
| Sol-4 | M | Sol | WT | 33.32154 | 18.6101 | 14.71143 | -1.88184 | 3.685438 |
| Sol-5 | F | Sol | WT | 31.93032 | 18.32911 | 13.60121 | -2.99206 | 7.956078 |
| Sol-6 | M | Sol | WT | 31.71953 | 18.06586 | 13.65367 | -2.9396 | 7.672004 |
| Sol-7 | F | Sol | G93A | 27.38946 | 19.44874 | 7.940718 | -8.65255 | 402.4182 |
| Sol-8 | F | Sol | G93A | 27.9512 | 18.92502 | 9.026186 | -7.56708 | 189.6353 |
| Sol-9 | M | Sol | G93A | 28.02578 | 18.73302 | 9.292762 | -7.30051 | 157.642 |
| Sol-10 | M | Sol | G93A | 28.28078 | 20.61494 | 7.665839 | -8.92743 | 486.8829 |
| Sol-11 | F | Sol | G93A | 25.41995 | 18.25901 | 7.160944 | -9.43233 | 690.8966 |
| Sol-12 | M | Sol | G93A | 27.11561 | 17.84087 | 9.274739 | -7.31853 | 159.6237 |
| Sol-13 | F | Sol | G93A_NaBu | 26.8996 | 18.25091 | 8.648691 | -7.94458 | 246.3522 |
| Sol-14 | M | Sol | G93A_NaBu | 31.41167 | 17.92598 | 13.48569 | -3.10758 | 8.619326 |
| Sol-15 | F | Sol | G93A_NaBu | 29.90428 | 19.34174 | 10.56253 | -6.03074 | 65.37808 |
| Sol-16 | F | Sol | G93A_NaBu | 27.95341 | 17.08681 | 10.8666 | -5.72667 | 52.95406 |
| Sol-17 | M | Sol | G93A_NaBu | 26.92578 | 17.92699 | 8.998786 | -7.59448 | 193.2713 |
| Dia-1 | F | Dia | WT | 30.45308 | 17.68709 | 12.76598 | -3.82729 | 14.19475 |
| Dia-2 | F | Dia | WT | 31.71014 | 18.08906 | 13.62108 | -2.97219 | 7.84727 |
| Dia-3 | M | Dia | WT | 31.42099 | 17.74711 | 13.67389 | -2.91938 | 7.565217 |
| Dia-4 | M | Dia | WT | 31.71808 | 17.6805 | 14.03758 | -2.55569 | 5.879466 |
| Dia-5 | F | Dia | WT | 30.94613 | 16.97426 | 13.97186 | -2.62141 | 6.153504 |
| Dia-6 | M | Dia | WT | 30.13928 | 16.97864 | 13.16064 | -3.43263 | 10.79754 |
| Dia-7 | F | Dia | G93A | 30.43077 | 20.90613 | 9.524638 | -7.06863 | 134.2363 |
| Dia-8 | F | Dia | G93A | 29.30744 | 20.77888 | 8.528563 | -8.06471 | 267.7433 |
| Dia-9 | M | Dia | G93A | 28.40634 | 17.72127 | 10.68508 | -5.90819 | 60.05424 |
| Dia-10 | M | Dia | G93A | 27.9058 | 17.83412 | 10.07167 | -6.5216 | 91.8747 |
| Dia-11 | F | Dia | G93A | 26.53009 | 17.41953 | 9.110562 | -7.48271 | 178.8626 |
| Dia-12 | M | Dia | G93A | 26.58765 | 16.4004 | 10.18725 | -6.40602 | 84.80162 |
| Dia-13 | F | Dia | G93A_NaBu | 27.88714 | 16.68548 | 11.20166 | -5.39161 | 41.97935 |
| Dia-14 | M | Dia | G93A_NaBu | 27.18891 | 16.68671 | 10.5022 | -6.09107 | 68.17036 |
| Dia-15 | F | Dia | G93A_NaBu | 31.99231 | 17.28896 | 14.70335 | -1.88992 | 3.706158 |
| Dia-16 | F | Dia | G93A_NaBu | 27.88591 | 17.48022 | 10.40569 | -6.18758 | 72.88633 |
| Dia-17 | M | Dia | G93A_NaBu | 27.01397 | 16.97117 | 10.0428 | -6.55047 | 93.73233 |
| Dia-18 | F | Dia | G93A_NaBu | 27.03133 | 16.20236 | 10.82896 | -5.76431 | 54.35368 |
| EOM-1 | F | EOM | WT | 31.87129 | 17.92036 | 13.95093 | -2.64234 | 6.243437 |
| EOM-2 | F | EOM | WT | 34.01603 | 17.79055 | 16.22548 | -0.36779 | 1.290373 |
| EOM-3 | M | EOM | WT | 32.97344 | 17.73169 | 15.24175 | -1.35152 | 2.551804 |
| EOM-4 | M | EOM | WT | 32.85031 | 16.79759 | 16.05272 | -0.54055 | 1.454523 |
| EOM-5 | F | EOM | WT | 32.14459 | 17.18029 | 14.9643 | -1.62897 | 3.092931 |
| EOM-6 | M | EOM | WT | 30.10094 | 16.2696 | 13.83135 | -2.76192 | 6.783004 |
| EOM-7 | F | EOM | G93A | 34.46942 | 20.38284 | 14.08658 | -2.50669 | 5.683146 |
| EOM-8 | F | EOM | G93A | 32.81526 | 18.65118 | 14.16408 | -2.42919 | 5.385901 |
| EOM-9 | M | EOM | G93A | 33.13782 | 17.68614 | 15.45168 | -1.14159 | 2.206244 |
| EOM-10 | M | EOM | G93A | 33.82894 | 18.21243 | 15.6165 | -0.97677 | 1.968048 |
| EOM-11 | F | EOM | G93A | 28.42253 | 16.55762 | 11.86491 | -4.72836 | 26.50813 |
| EOM-12 | M | EOM | G93A | 30.90527 | 17.52344 | 13.38183 | -3.21144 | 9.262745 |
| EOM-13 | F | EOM | G93A_NaBu | 30.2457 | 15.9794 | 14.2663 | -2.32697 | 5.017514 |
| EOM-14 | M | EOM | G93A_NaBu | 29.83213 | 15.87309 | 13.95904 | -2.63423 | 6.208446 |
| EOM-15 | M | EOM | G93A_NaBu | 29.8665 | 16.07902 | 13.78748 | -2.80579 | 6.992414 |
| EOM-16 | F | EOM | G93A_NaBu | 29.88214 | 15.80923 | 14.07291 | -2.52036 | 5.737263 |
| EOM-17 | M | EOM | G93A_NaBu | 31.03522 | 18.17189 | 12.86333 | -3.72994 | 13.26855 |
| EOM-18 | M | EOM | G93A_NaBu | 30.50419 | 16.9355 | 13.56869 | -3.02458 | 8.137472 |
