## Supplementary material for "Distinct transcriptomic profile of satellite cells contributes to preservation of neuromuscular junctions in extraocular muscles of ALS mice": Figure 2-Source Data 1

**Figure 2-Source Data 1. Quantification results of averaged peri-NMJ SC numbers in muscles of different origins and treatment conditions.**

| Mouse | Gender | Muscle | Group | Mean SC No at NMJ | NMJ counted |
| --- | --- | --- | --- | --- | --- |
| EDL-1 | M | EDL | WT | 0.47619 | 63 |
| EDL-2 | M | EDL | WT | 0.4375 | 48 |
| EDL-3 | F | EDL | WT | 0.395833333 | 48 |
| EDL-4 | F | EDL | WT | 0.460526316 | 76 |
| EDL-5 | M | EDL | G93A | 0.064748201 | 139 |
| EDL-6 | M | EDL | G93A | 0.063492063 | 63 |
| EDL-7 | F | EDL | G93A | 0.014492754 | 69 |
| EDL-8 | F | EDL | G93A | 0.114285714 | 35 |
| EDL-9 | M | EDL | G93A | 0.185714286 | 70 |
| EDL-10 | F | EDL | G93A | 0.184615385 | 65 |
| EDL-11 | F | EDL | G93A_NaBu | 0.481481481 | 37 |
| EDL-12 | M | EDL | G93A_NaBu | 0.166666667 | 48 |
| EDL-13 | M | EDL | G93A_NaBu | 0.357142857 | 28 |
| EDL-14 | F | EDL | G93A_NaBu | 0.355555556 | 45 |
| EDL-15 | F | EDL | G93A_NaBu | 0.162162162 | 37 |
| EDL-16 | F | EDL | G93A_NaBu | 0.325581395 | 43 |
| Sol-1 | M | Sol | WT | 3.05 | 20 |
| Sol-2 | F | Sol | WT | 2.875 | 24 |
| Sol-3 | F | Sol | WT | 2.75 | 12 |
| Sol-4 | F | Sol | WT | 3.071428571 | 42 |
| Sol-5 | M | Sol | WT | 3 | 45 |
| Sol-6 | M | Sol | G93A | 1.139534884 | 43 |
| Sol-7 | M | Sol | G93A | 1.357142857 | 14 |
| Sol-8 | M | Sol | G93A | 0.923076923 | 26 |
| Sol-9 | F | Sol | G93A | 0.552631579 | 38 |
| Sol-10 | F | Sol | G93A | 0.756097561 | 41 |
| Sol-11 | F | Sol | G93A_NaBu | 3 | 12 |
| Sol-12 | M | Sol | G93A_NaBu | 3.166666667 | 12 |
| Sol-13 | F | Sol | G93A_NaBu | 0.888888889 | 18 |
| Sol-14 | F | Sol | G93A_NaBu | 2 | 8 |
| Sol-15 | F | Sol | G93A_NaBu | 3.428571429 | 14 |
| Dia-1 | M | Dia | WT | 2.983333 | 120 |
| Dia-2 | F | Dia | WT | 1.987179487 | 78 |
| Dia-3 | F | Dia | WT | 2.32 | 25 |
| Dia-4 | F | Dia | WT | 2.882352941 | 51 |
| Dia-5 | F | Dia | WT | 2.830357143 | 112 |
| Dia-6 | M | Dia | G93A | 2 | 36 |
| Dia-7 | F | Dia | G93A | 2.4 | 70 |
| Dia-8 | F | Dia | G93A | 1.04 | 50 |
| Dia-9 | F | Dia | G93A | 1.188888889 | 90 |
| Dia-10 | F | Dia | G93A_NaBu | 2.095238095 | 42 |
| Dia-11 | F | Dia | G93A_NaBu | 2.795454545 | 44 |
| Dia-12 | F | Dia | G93A_NaBu | 3.21875 | 64 |
| Dia-13 | F | Dia | G93A_NaBu | 2.693333333 | 75 |
| Dia-14 | F | Dia | G93A | 1.47541 | 183 |
| EOM-1 | M | EOM | WT | 3.22807 | 114 |
| EOM-2 | F | EOM | WT | 2.965517241 | 29 |
| EOM-3 | F | EOM | WT | 3.222222222 | 9 |
| EOM-4 | F | EOM | WT | 3.081632653 | 49 |
| EOM-5 | F | EOM | G93A | 2.375 | 8 |
| EOM-6 | F | EOM | G93A | 2.507692308 | 65 |
| EOM-7 | F | EOM | G93A | 2.451612903 | 31 |
| EOM-8 | F | EOM | G93A | 3.285714 | 105 |
| EOM-9 | F | EOM | G93A | 3.142857 | 126 |
| EOM-10 | M | EOM | G93A_NaBu | 2.642105 | 95 |
| EOM-11 | F | EOM | G93A_NaBu | 2.680851064 | 47 |
| EOM-12 | F | EOM | G93A_NaBu | 2.329545455 | 88 |
| EOM-13 | F | EOM | G93A_NaBu | 2.730769231 | 26 |
