## Supplementary material for "Distinct transcriptomic profile of satellite cells contributes to preservation of neuromuscular junctions in extraocular muscles of ALS mice": Figure 3-Source Data 1

**Figure 3-Source Data 1. Percentage of P5-4 events recorded in different rounds of sorting.**

| Batch ID | Gender | Muscle | Group | P5-4 percentage |
| --- | --- | --- | --- | --- |
| HL-1 | M | HL | WT | 8.03 |
| HL-2 | M | HL | WT | 7.12 |
| HL-3 | F | HL | WT | 9.31 |
| HL-4 | M | HL | WT | 6.59 |
| HL-5 | F | HL | WT | 5.85 |
| HL-6 | F | HL | WT | 6.4 |
| HL-7 | F | HL | WT | 7.64 |
| HL-8 | F | HL | WT | 7.42 |
| HL-9 | F | HL | WT | 6.25 |
| HL-10 | M | HL | WT | 11.12 |
| HL-11 | M | HL | WT | 8.39 |
| HL-12 | M | HL | WT | 8.39 |
| HL-13 | M | HL | WT | 8.08 |
| HL-14 | M | HL | WT | 7.33 |
| HL-15 | F | HL | G93A | 7.64 |
| HL-16 | M | HL | G93A | 3.72 |
| HL-17 | M | HL | G93A | 5.29 |
| HL-18 | F | HL | G93A | 5.29 |
| HL-19 | M | HL | G93A | 2.45 |
| HL-20 | M | HL | G93A | 8.81 |
| HL-21 | F | HL | G93A | 4 |
| HL-22 | F | HL | G93A | 4.17 |
| HL-23 | M | HL | G93A | 3.45 |
| HL-24 | M | HL | G93A | 5.78 |
| HL-25 | F | HL | G93A | 6.93 |
| HL-26 | M | HL | G93A | 5.59 |
| HL-27 | M | HL | G93A | 5.7 |
| HL-28 | M | HL | G93A | 5.45 |
| Dia-1 | F | Dia | WT | 13.03 |
| Dia-2 | M | Dia | WT | 10.7 |
| Dia-3 | M | Dia | WT | 9.91 |
| Dia-4 | F | Dia | WT | 11.85 |
| Dia-5 | F | Dia | WT | 15.73 |
| Dia-6 | M | Dia | WT | 13.47 |
| Dia-7 | M | Dia | WT | 10.99 |
| Dia-8 | M | Dia | WT | 18.19 |
| Dia-9 | M | Dia | WT | 12.77 |
| Dia-10 | M | Dia | G93A | 8.06 |
| Dia-11 | F | Dia | G93A | 9.41 |
| Dia-12 | M | Dia | G93A | 3.06 |
| Dia-13 | M | Dia | G93A | 0.91 |
| Dia-14 | F | Dia | G93A | 16.8 |
| Dia-15 | F | Dia | G93A | 6.45 |
| Dia-16 | M | Dia | G93A | 9.53 |
| Dia-17 | M | Dia | G93A | 13.04 |
| Dia-18 | F | Dia | G93A | 7.56 |
| Dia-19 | M | Dia | G93A | 8.92 |
| Dia-20 | M | Dia | G93A | 8.5 |
| Dia-21 | F | Dia | G93A | 15.83 |
| Dia-22 | M | Dia | G93A | 11.01 |
| Dia-23 | M | Dia | G93A | 15.91 |
| Dia-24 | M | Dia | G93A | 13.53 |
| Dia-25 | M | Dia | G93A | 11.23 |
| Dia-26 | M | Dia | G93A | 11.91 |
| EOM-1 | F | EOM | WT | 7.89 |
| EOM-2 | M | EOM | WT | 7.61 |
| EOM-3 | M | EOM | WT | 10.28 |
| EOM-4 | M | EOM | WT | 9.51 |
| EOM-5 | M | EOM | WT | 10.35 |
| EOM-6 | M | EOM | WT | 12.18 |
| EOM-7 | M | EOM | WT | 12.21 |
| EOM-8 | M | EOM | G93A | 3.29 |
| EOM-9 | F | EOM | G93A | 3.72 |
| EOM-10 | F | EOM | G93A | 9.71 |
| EOM-11 | M | EOM | G93A | 9 |
| EOM-12 | M | EOM | G93A | 9.5 |
| EOM-13 | M | EOM | G93A | 9.7 |
| EOM-14 | M | EOM | G93A | 8.16 |
