## Supplementary material for "Distinct transcriptomic profile of satellite cells contributes to preservation of neuromuscular junctions in extraocular muscles of ALS mice": Figure 6-Source Data 1

**Figure 6-Source Data 1. Top 20 differentially expressed genes comparing EOM SCs to hindlimb and diaphragm counterparts cultured in growth and differentiation medium.** Log2FC and FDR are shown and genes are ranked according to log2FC. Column 2-5 are genes expressed higher in EOM SCs and Column 6-9 are genes expressed lower in EOM SCs.

| Group of comparison | Gene ID (higher) | Gene Symbol | log2 | FDR | Gene ID (lower) | Gene Symbol | log2 | FDR |
| --- | --- | --- | --- | --- | --- | --- | --- | --- |
| WT_EOM_G/WT_HL_G | 11695 | 'Alx4' | 10.82496 | 0 | 16814 | 'Lbx1' | -10.2825 | 0 |
| WT_EOM_G/WT_HL_G | 111241 | 'Hmga1b' | 9.440437 | 0 | 15427 | 'Hoxc9' | -10.0196 | 5.44E-90 |
| WT_EOM_G/WT_HL_G | 18740 | 'Pitx1' | 9.019591 | 1.11E-63 | 15077 | 'Hist2h3c1' | -9.30834 | 1.31E-19 |
| WT_EOM_G/WT_HL_G | 22441 | 'Xlr' | 8.022368 | 5.14E-14 | 15404 | 'Hoxa7' | -9.28309 | 1.44E-61 |
| WT_EOM_G/WT_HL_G | 16371 | 'Irx1' | 7.741467 | 2.80E-22 | 15425 | 'Hoxc6' | -8.5727 | 1.63E-246 |
| WT_EOM_G/WT_HL_G | 102633156 | 'LOC102633156' | 7.434628 | 1.56E-36 | 15402 | 'Hoxa5' | -8.56605 | 6.28E-36 |
| WT_EOM_G/WT_HL_G | 105827 | 'Amigo2' | 6.442943 | 1.01E-13 | 15401 | 'Hoxa4' | -8.20945 | 1.44E-22 |
| WT_EOM_G/WT_HL_G | 13076 | 'Cyp1a1' | 6.253341 | 2.65E-66 | 15400 | 'Hoxa3' | -7.83289 | 1.09E-40 |
| WT_EOM_G/WT_HL_G | 219257 | 'Pcdh20' | 6.022368 | 7.21E-48 | 15426 | 'Hoxc8' | -7.54689 | 1.44E-22 |
| WT_EOM_G/WT_HL_G | 216616 | 'Efemp1' | 5.785102 | 7.42E-34 | 15403 | 'Hoxa6' | -7.17991 | 5.47E-05 |
| WT_EOM_G/WT_HL_G | 242620 | 'Dmrta2' | 5.777608 | 1.39E-214 | 15394 | 'Hoxa1' | -5.79181 | 8.74E-30 |
| WT_EOM_G/WT_HL_G | 17389 | 'Mmp16' | 5.643856 | 5.50E-33 | 15424 | 'Hoxc5' | -5.7211 | 2.54E-26 |
| WT_EOM_G/WT_HL_G | 14049 | 'Eya2' | 5.566477 | 5.03E-183 | 18505 | 'Pax3' | -5.29768 | 1.38E-18 |
| WT_EOM_G/WT_HL_G | 110648 | 'Lmx1a' | 5.426265 | 1.41E-07 | 268527 | 'Greb1' | -5.17136 | 0 |
| WT_EOM_G/WT_HL_G | 20257 | 'Stmn2' | 5.373458 | 1.14E-265 | 13628 | 'Eef1a2' | -5.03148 | 1.27E-48 |
| WT_EOM_G/WT_HL_G | 70691 | '3830403N18Rik' | 5.254241 | 4.25E-10 | 100702 | 'Gbp6' | -4.65372 | 0 |
| WT_EOM_G/WT_HL_G | 20274 | 'Scn9a' | 5.111508 | 1.04E-98 | 12740 | 'Cldn4' | -4.60554 | 8.89E-52 |
| WT_EOM_G/WT_HL_G | 14402 | 'Gabrb3' | 5.044394 | 1.28E-09 | 21380 | 'Tbx1' | -4.52061 | 4.03E-143 |
| WT_EOM_G/WT_HL_G | 17286 | 'Meox2' | 5.022368 | 9.37E-07 | 76074 | 'Gbp8' | -4.40273 | 0 |
| WT_EOM_G/WT_HL_G | 12824 | 'Col2a1' | 4.922832 | 2.68E-20 | 626578 | 'Gbp10' | -4.33273 | 0 |
| WT_EOM_G/WT_Dia_G | 18740 | 'Pitx1' | 9.019591 | 1.18E-63 | 15402 | 'Hoxa5' | -10.5498 | 1.14E-142 |
| WT_EOM_G/WT_Dia_G | 11695 | 'Alx4' | 8.239996 | 0 | 15401 | 'Hoxa4' | -10.4798 | 2.12E-109 |
| WT_EOM_G/WT_Dia_G | 219257 | 'Pcdh20' | 7.60733 | 6.03E-51 | 16814 | 'Lbx1' | -10.0014 | 2.73E-270 |
| WT_EOM_G/WT_Dia_G | 20274 | 'Scn9a' | 6.918863 | 7.93E-110 | 15400 | 'Hoxa3' | -9.85175 | 2.26E-139 |
| WT_EOM_G/WT_Dia_G | 17389 | 'Mmp16' | 6.643856 | 2.15E-34 | 15404 | 'Hoxa7' | -9.32418 | 2.74E-63 |
| WT_EOM_G/WT_Dia_G | 12824 | 'Col2a1' | 6.507795 | 2.85E-24 | 15077 | 'Hist2h3c1' | -8.90989 | 8.28E-15 |
| WT_EOM_G/WT_Dia_G | 54519 | 'Apbb1ip' | 6.303781 | 8.21E-29 | 15425 | 'Hoxc6' | -8.82972 | 4.24E-176 |
| WT_EOM_G/WT_Dia_G | 13076 | 'Cyp1a1' | 6.253341 | 2.78E-66 | 18505 | 'Pax3' | -7.78354 | 7.37E-119 |
| WT_EOM_G/WT_Dia_G | 236285 | 'Lancl3' | 6.228819 | 1.01E-13 | 15427 | 'Hoxc9' | -7.45121 | 4.21E-15 |
| WT_EOM_G/WT_Dia_G | 16371 | 'Irx1' | 6.156504 | 1.23E-20 | 15426 | 'Hoxc8' | -7.07682 | 2.81E-16 |
| WT_EOM_G/WT_Dia_G | 16917 | 'Lmx1b' | 6.087463 | 1.14E-25 | 15394 | 'Hoxa1' | -6.97499 | 1.82E-70 |
| WT_EOM_G/WT_Dia_G | 17286 | 'Meox2' | 6.022368 | 8.32E-08 | 15403 | 'Hoxa6' | -6.88264 | 4.48E-04 |
| WT_EOM_G/WT_Dia_G | 19126 | 'Prom1' | 6.022368 | 2.67E-11 | 15424 | 'Hoxc5' | -6.87036 | 1.31E-61 |
| WT_EOM_G/WT_Dia_G | 12296 | 'Cacnb2' | 6.022368 | 6.19E-28 | 21380 | 'Tbx1' | -5.05004 | 1.61E-220 |
| WT_EOM_G/WT_Dia_G | 13134 | 'Dach1' | 5.906891 | 2.70E-16 | 15423 | 'Hoxc4' | -4.87228 | 1.25E-117 |
| WT_EOM_G/WT_Dia_G | 58198 | 'Sall1' | 5.72792 | 3.09E-14 | 100702 | 'Gbp6' | -4.37603 | 0 |
| WT_EOM_G/WT_Dia_G | 242620 | 'Dmrta2' | 5.4975 | 8.82E-212 | 13628 | 'Eef1a2' | -4.36375 | 2.92E-28 |
| WT_EOM_G/WT_Dia_G | 22441 | 'Xlr' | 5.437405 | 1.40E-12 | 626578 | 'Gbp10' | -4.12044 | 0 |
| WT_EOM_G/WT_Dia_G | 71690 | 'Esm1' | 5.357552 | 4.45E-23 | 76074 | 'Gbp8' | -4.07852 | 0 |
| WT_EOM_G/WT_Dia_G | 12737 | 'Cldn1' | 5.357552 | 6.16E-07 | 65255 | 'Asb4' | -3.65208 | 4.77E-21 |
| G93A_EOM_G/G93A_HL_G | 11695 | 'Alx4' | 10.05393 | 0 | 15404 | 'Hoxa7' | -8.89785 | 1.23E-46 |
| G93A_EOM_G/G93A_HL_G | 18740 | 'Pitx1' | 9.266787 | 1.14E-75 | 15077 | 'Hist2h3c1' | -8.55075 | 2.43E-11 |
| G93A_EOM_G/G93A_HL_G | 22441 | 'Xlr' | 7.924813 | 4.04E-12 | 15402 | 'Hoxa5' | -8.34873 | 1.39E-30 |
| G93A_EOM_G/G93A_HL_G | 70691 | '3830403N18Rik' | 7.285402 | 8.30E-08 | 15425 | 'Hoxc6' | -8.31628 | 3.64E-247 |
| G93A_EOM_G/G93A_HL_G | 17389 | 'Mmp16' | 6.285402 | 1.85E-28 | 15401 | 'Hoxa4' | -8.14975 | 1.56E-21 |
| G93A_EOM_G/G93A_HL_G | 219257 | 'Pcdh20' | 6.273018 | 7.47E-57 | 15427 | 'Hoxc9' | -8.13955 | 2.14E-70 |
| G93A_EOM_G/G93A_HL_G | 242620 | 'Dmrta2' | 6.112391 | 3.34E-258 | 15426 | 'Hoxc8' | -7.89482 | 1.81E-28 |
| G93A_EOM_G/G93A_HL_G | 12296 | 'Cacnb2' | 5.97728 | 3.48E-14 | 15403 | 'Hoxa6' | -7.55459 | 2.67E-06 |
| G93A_EOM_G/G93A_HL_G | 16371 | 'Irx1' | 5.83289 | 3.65E-16 | 15400 | 'Hoxa3' | -7.36632 | 5.56E-30 |
| G93A_EOM_G/G93A_HL_G | 13076 | 'Cyp1a1' | 5.264107 | 7.04E-31 | 15394 | 'Hoxa1' | -7.26053 | 2.86E-52 |
| G93A_EOM_G/G93A_HL_G | 110648 | 'Lmx1a' | 5.247928 | 1.18E-11 | 15424 | 'Hoxc5' | -6.48382 | 3.05E-23 |
| G93A_EOM_G/G93A_HL_G | 102633156 | 'LOC102633156' | 5.209453 | 8.30E-08 | 16814 | 'Lbx1' | -6.06757 | 1.30E-256 |
| G93A_EOM_G/G93A_HL_G | 22160 | 'Twist1' | 5.129283 | 3.85E-36 | 21380 | 'Tbx1' | -5.67304 | 2.03E-190 |
| G93A_EOM_G/G93A_HL_G | 20274 | 'Scn9a' | 5.053111 | 7.09E-72 | 12740 | 'Cldn4' | -4.87893 | 7.23E-73 |
| G93A_EOM_G/G93A_HL_G | 57246 | 'Tbx20' | 5.044394 | 1.14E-08 | 626578 | 'Gbp10' | -4.85039 | 2.94E-198 |
| G93A_EOM_G/G93A_HL_G | 20257 | 'Stmn2' | 5.030934 | 0 | 18505 | 'Pax3' | -4.52356 | 1.66E-07 |
| G93A_EOM_G/G93A_HL_G | 14049 | 'Eya2' | 5.015202 | 1.62E-188 | 76074 | 'Gbp8' | -4.48187 | 3.00E-228 |
| G93A_EOM_G/G93A_HL_G | 16917 | 'Lmx1b' | 5.008989 | 2.09E-35 | 100702 | 'Gbp6' | -4.25481 | 0 |
| G93A_EOM_G/G93A_HL_G | 58198 | 'Sall1' | 4.930737 | 1.31E-13 | 99738 | 'Kcnc4' | -4.10852 | 3.15E-11 |
| G93A_EOM_G/G93A_HL_G | 71690 | 'Esm1' | 4.890771 | 2.48E-08 | 22771 | 'Zic1' | -3.97982 | 5.10E-19 |
| G93A_EOM_G/G93A_Dia_G | 111241 | 'Hmga1b' | 13.81898 | 0 | 15401 | 'Hoxa4' | -10.5459 | 5.03E-114 |
| G93A_EOM_G/G93A_Dia_G | 18740 | 'Pitx1' | 9.266787 | 8.08E-77 | 15402 | 'Hoxa5' | -10.5018 | 1.94E-137 |
| G93A_EOM_G/G93A_Dia_G | 22441 | 'Xlr' | 7.924813 | 3.43E-12 | 15400 | 'Hoxa3' | -10.1434 | 2.34E-168 |
| G93A_EOM_G/G93A_Dia_G | 11695 | 'Alx4' | 7.731998 | 0 | 15425 | 'Hoxc6' | -9.01053 | 1.78E-208 |
| G93A_EOM_G/G93A_Dia_G | 16371 | 'Irx1' | 7.417853 | 8.73E-18 | 15404 | 'Hoxa7' | -8.87344 | 4.65E-46 |
| G93A_EOM_G/G93A_Dia_G | 219257 | 'Pcdh20' | 6.857981 | 2.52E-59 | 15077 | 'Hist2h3c1' | -8.13955 | 1.08E-08 |
| G93A_EOM_G/G93A_Dia_G | 20257 | 'Stmn2' | 6.430542 | 0 | 18505 | 'Pax3' | -8.13955 | 3.55E-105 |
| G93A_EOM_G/G93A_Dia_G | 110648 | 'Lmx1a' | 6.247928 | 4.53E-13 | 15424 | 'Hoxc5' | -7.96 | 2.56E-67 |
| G93A_EOM_G/G93A_Dia_G | 13134 | 'Dach1' | 6.087463 | 1.79E-17 | 15394 | 'Hoxa1' | -7.80735 | 4.98E-78 |
| G93A_EOM_G/G93A_Dia_G | 14049 | 'Eya2' | 6.039864 | 4.45E-208 | 15426 | 'Hoxc8' | -7.30378 | 5.64E-19 |
| G93A_EOM_G/G93A_Dia_G | 11830 | 'Aqp5' | 5.931915 | 0 | 15403 | 'Hoxa6' | -7.19967 | 5.89E-05 |
| G93A_EOM_G/G93A_Dia_G | 58198 | 'Sall1' | 5.930737 | 2.61E-16 | 21380 | 'Tbx1' | -5.91783 | 9.52E-231 |
| G93A_EOM_G/G93A_Dia_G | 17286 | 'Meox2' | 5.906891 | 3.06E-07 | 15427 | 'Hoxc9' | -5.72792 | 2.60E-12 |
| G93A_EOM_G/G93A_Dia_G | 19126 | 'Prom1' | 5.61471 | 2.85E-09 | 16814 | 'Lbx1' | -5.70617 | 1.73E-197 |
| G93A_EOM_G/G93A_Dia_G | 208898 | 'Unc13c' | 5.491853 | 2.46E-18 | 626578 | 'Gbp10' | -5.67181 | 0 |
| G93A_EOM_G/G93A_Dia_G | 20274 | 'Scn9a' | 5.375039 | 4.20E-73 | 76074 | 'Gbp8' | -5.64684 | 0 |
| G93A_EOM_G/G93A_Dia_G | 242620 | 'Dmrta2' | 5.282316 | 3.88E-247 | 65255 | 'Asb4' | -4.81762 | 2.23E-21 |
| G93A_EOM_G/G93A_Dia_G | 13076 | 'Cyp1a1' | 5.264107 | 2.42E-31 | 228801 | 'Bpifb1' | -4.40939 | 3.61E-06 |
| G93A_EOM_G/G93A_Dia_G | 18573 | 'Pde1a' | 5.220551 | 1.84E-34 | 100702 | 'Gbp6' | -4.34405 | 0 |
| G93A_EOM_G/G93A_Dia_G | 54352 | 'Irx5' | 5.147754 | 5.68E-188 | 68701 | 'Ppp1r27' | -4.33213 | 1.25E-87 |
| WT_EOM_D/WT_HL_D | 111241 | 'Hmga1b' | 9.340221 | 0 | 209448 | 'Hoxc10' | -12 | 0 |
| WT_EOM_D/WT_HL_D | 14699 | 'Gngt1' | 8.475733 | 2.17E-16 | 15405 | 'Hoxa9' | -9.57175 | 1.03E-123 |
| WT_EOM_D/WT_HL_D | 18740 | 'Pitx1' | 8.312883 | 6.01E-38 | 15427 | 'Hoxc9' | -9.55267 | 1.18E-64 |
| WT_EOM_D/WT_HL_D | 70691 | '3830403N18Rik' | 7.651052 | 6.46E-10 | 15404 | 'Hoxa7' | -7.84967 | 4.63E-66 |
| WT_EOM_D/WT_HL_D | 20272 | 'Scn7a' | 7.169925 | 1.61E-102 | 15426 | 'Hoxc8' | -7.70736 | 3.42E-25 |
| WT_EOM_D/WT_HL_D | 14049 | 'Eya2' | 7.014578 | 8.64E-143 | 15425 | 'Hoxc6' | -6.92103 | 8.27E-202 |
| WT_EOM_D/WT_HL_D | 16371 | 'Irx1' | 6.564785 | 3.59E-27 | 13628 | 'Eef1a2' | -6.16993 | 7.99E-21 |
| WT_EOM_D/WT_HL_D | 94332 | 'Cadm3' | 6.426265 | 3.79E-22 | 21380 | 'Tbx1' | -5.91049 | 3.96E-100 |
| WT_EOM_D/WT_HL_D | 11695 | 'Alx4' | 6.322443 | 0 | 15402 | 'Hoxa5' | -5.73245 | 4.22E-27 |
| WT_EOM_D/WT_HL_D | 17389 | 'Mmp16' | 6.066089 | 3.34E-20 | 15401 | 'Hoxa4' | -5.53657 | 1.24E-37 |
| WT_EOM_D/WT_HL_D | 74318 | 'Hopx' | 5.964278 | 6.09E-70 | 15399 | 'Hoxa2' | -5.04439 | 7.12E-08 |
| WT_EOM_D/WT_HL_D | 216616 | 'Efemp1' | 5.924813 | 2.18E-84 | 207215 | 'Fbxo40' | -4.94854 | 1.24E-228 |
| WT_EOM_D/WT_HL_D | 20257 | 'Stmn2' | 5.855108 | 4.76E-126 | 545611 | 'Fam205a2' | -4.85798 | 2.77E-06 |
| WT_EOM_D/WT_HL_D | 19263 | 'Ptprb' | 5.83289 | 2.63E-31 | 241431 | 'Xirp2' | -4.57218 | 0 |
| WT_EOM_D/WT_HL_D | 12023 | 'Barx2' | 5.784635 | 1.59E-103 | 15400 | 'Hoxa3' | -4.54844 | 9.35E-21 |
| WT_EOM_D/WT_HL_D | 16008 | 'Igfbp2' | 5.682558 | 0 | 15394 | 'Hoxa1' | -4.35755 | 3.33E-19 |
| WT_EOM_D/WT_HL_D | 15214 | 'Hey2' | 5.584963 | 2.77E-06 | 16814 | 'Lbx1' | -4.28221 | 1.35E-204 |
| WT_EOM_D/WT_HL_D | 15957 | 'Ifit1' | 5.57289 | 5.48E-28 | 15424 | 'Hoxc5' | -4.18982 | 9.47E-16 |
| WT_EOM_D/WT_HL_D | 109323 | 'C1qtnf7' | 5.392317 | 1.27E-04 | 68701 | 'Ppp1r27' | -4.09359 | 5.12E-70 |
| WT_EOM_D/WT_HL_D | 110648 | 'Lmx1a' | 5.357552 | 4.05E-07 | 235416 | 'Lman1l' | -4.00956 | 3.43E-48 |
| WT_EOM_D/WT_Dia_D | 14699 | 'Gngt1' | 8.475733 | 1.54E-16 | 15427 | 'Hoxc9' | -8.21917 | 8.32E-26 |
| WT_EOM_D/WT_Dia_D | 18740 | 'Pitx1' | 8.312883 | 2.52E-38 | 15425 | 'Hoxc6' | -7.85968 | 3.33E-233 |
| WT_EOM_D/WT_Dia_D | 20272 | 'Scn7a' | 8.169925 | 1.39E-105 | 15404 | 'Hoxa7' | -7.82231 | 5.66E-65 |
| WT_EOM_D/WT_Dia_D | 14049 | 'Eya2' | 8.014578 | 4.06E-148 | 15426 | 'Hoxc8' | -7.73471 | 8.32E-26 |
| WT_EOM_D/WT_Dia_D | 74318 | 'Hopx' | 7.902878 | 1.07E-75 | 15402 | 'Hoxa5' | -7.47438 | 4.73E-97 |
| WT_EOM_D/WT_Dia_D | 20319 | 'Sfrp2' | 6.77259 | 2.01E-29 | 21380 | 'Tbx1' | -6.66747 | 3.13E-176 |
| WT_EOM_D/WT_Dia_D | 216616 | 'Efemp1' | 6.602884 | 1.93E-88 | 13628 | 'Eef1a2' | -6.47032 | 5.28E-26 |
| WT_EOM_D/WT_Dia_D | 20257 | 'Stmn2' | 6.555548 | 4.01E-132 | 545611 | 'Fam205a2' | -5.70044 | 9.39E-11 |
| WT_EOM_D/WT_Dia_D | 20855 | 'Stc1' | 6.507795 | 2.15E-17 | 15399 | 'Hoxa2' | -5.60486 | 7.28E-12 |
| WT_EOM_D/WT_Dia_D | 11695 | 'Alx4' | 6.129798 | 0 | 15400 | 'Hoxa3' | -5.57894 | 3.47E-35 |
| WT_EOM_D/WT_Dia_D | 17389 | 'Mmp16' | 6.066089 | 5.80E-22 | 15401 | 'Hoxa4' | -5.51307 | 3.78E-37 |
| WT_EOM_D/WT_Dia_D | 242620 | 'Dmrta2' | 5.860742 | 4.53E-211 | 209448 | 'Hoxc10' | -5.39232 | 2.11E-04 |
| WT_EOM_D/WT_Dia_D | 16775 | 'Lama4' | 5.857981 | 5.80E-18 | 15424 | 'Hoxc5' | -5.27146 | 4.30E-37 |
| WT_EOM_D/WT_Dia_D | 12819 | 'Col15a1' | 5.792248 | 1.75E-154 | 15405 | 'Hoxa9' | -4.90689 | 2.67E-05 |
| WT_EOM_D/WT_Dia_D | 26564 | 'Ror2' | 5.681824 | 1.01E-27 | 15394 | 'Hoxa1' | -4.83289 | 1.64E-28 |
| WT_EOM_D/WT_Dia_D | 11830 | 'Aqp5' | 5.563283 | 2.47E-278 | 140781 | 'Myh7' | -4.57388 | 0 |
| WT_EOM_D/WT_Dia_D | 12023 | 'Barx2' | 5.562242 | 2.55E-103 | 16814 | 'Lbx1' | -4.47908 | 8.89E-274 |
| WT_EOM_D/WT_Dia_D | 12835 | 'Col6a3' | 5.52147 | 0 | 207215 | 'Fbxo40' | -4.47524 | 2.46E-154 |
| WT_EOM_D/WT_Dia_D | 109323 | 'C1qtnf7' | 5.392317 | 1.16E-04 | 241431 | 'Xirp2' | -4.37851 | 0 |
| WT_EOM_D/WT_Dia_D | 16008 | 'Igfbp2' | 5.387391 | 0 | 15423 | 'Hoxc4' | -4.26595 | 9.63E-105 |
| G93A_EOM_D/G93A_HL_D | 12159 | 'Bmp4' | 9.355351 | 1.10E-57 | 209448 | 'Hoxc10' | 0 | -5.39232 |
| G93A_EOM_D/G93A_HL_D | 14049 | 'Eya2' | 8.423466 | 8.78E-121 | 15427 | 'Hoxc9' | 9.84E-69 | -8.21917 |
| G93A_EOM_D/G93A_HL_D | 18740 | 'Pitx1' | 8.055282 | 4.90E-31 | 15405 | 'Hoxa9' | 9.41E-116 | -4.90689 |
| G93A_EOM_D/G93A_HL_D | 20272 | 'Scn7a' | 7.684164 | 6.91E-218 | 15426 | 'Hoxc8' | 7.91E-18 | -7.73471 |
| G93A_EOM_D/G93A_HL_D | 12814 | 'Col11a1' | 7.366322 | 5.70E-58 | 15404 | 'Hoxa7' | 9.99E-50 | -7.82231 |
| G93A_EOM_D/G93A_HL_D | 70691 | '3830403N18Rik' | 6.906891 | 5.23E-06 | 56222 | 'Cited4' | 4.15E-05 | -4.12928 |
| G93A_EOM_D/G93A_HL_D | 22160 | 'Twist1' | 6.405141 | 7.29E-25 | 16814 | 'Lbx1' | 0 | -4.47908 |
| G93A_EOM_D/G93A_HL_D | 11695 | 'Alx4' | 6.359104 | 0 | 21380 | 'Tbx1' | 9.95E-109 | -6.66747 |
| G93A_EOM_D/G93A_HL_D | 20855 | 'Stc1' | 6.321928 | 1.26E-40 | 545611 | 'Fam205a2' | 6.40E-11 | -5.70044 |
| G93A_EOM_D/G93A_HL_D | 216616 | 'Efemp1' | 6.157541 | 8.96E-81 | 15402 | 'Hoxa5' | 3.19E-24 | -7.47438 |
| G93A_EOM_D/G93A_HL_D | 17389 | 'Mmp16' | 6.149747 | 8.13E-25 | 15425 | 'Hoxc6' | 6.80E-154 | -7.85968 |
| G93A_EOM_D/G93A_HL_D | 72107 | 'Dscc1' | 6.101538 | 9.79E-13 | 13628 | 'Eef1a2' | 3.21E-11 | -6.47032 |
| G93A_EOM_D/G93A_HL_D | 12023 | 'Barx2' | 5.848474 | 2.12E-93 | 15401 | 'Hoxa4' | 6.84E-21 | -5.51307 |
| G93A_EOM_D/G93A_HL_D | 16775 | 'Lama4' | 5.807355 | 1.82E-15 | 15424 | 'Hoxc5' | 1.34E-14 | -5.27146 |
| G93A_EOM_D/G93A_HL_D | 102371 | 'Myzap' | 5.798234 | 5.91E-94 | 15399 | 'Hoxa2' | 1.80E-06 | -5.60486 |
| G93A_EOM_D/G93A_HL_D | 56533 | 'Rgs17' | 5.78136 | 6.65E-41 | 14537 | 'Gcnt1' | 2.20E-16 | -3.42253 |
| G93A_EOM_D/G93A_HL_D | 12819 | 'Col15a1' | 5.609794 | 5.95E-68 | 15394 | 'Hoxa1' | 8.10E-26 | -4.83289 |
| G93A_EOM_D/G93A_HL_D | 434215 | 'Lrrc32' | 5.551796 | 4.01E-88 | 241431 | 'Xirp2' | 0 | -4.37851 |
| G93A_EOM_D/G93A_HL_D | 217843 | 'Unc79' | 5.523562 | 1.87E-19 | 17700 | 'Mstn' | 6.17E-102 | -2.13386 |
| G93A_EOM_D/G93A_HL_D | 12835 | 'Col6a3' | 5.510154 | 0 | 15400 | 'Hoxa3' | 7.00E-18 | -5.57894 |
| G93A_EOM_D/G93A_Dia_D | 111241 | 'Hmga1b' | 13.11943 | 0 | 15427 | 'Hoxc9' | -8.73132 | 1.39E-36 |
| G93A_EOM_D/G93A_Dia_D | 20272 | 'Scn7a' | 8.269127 | 5.37E-226 | 15426 | 'Hoxc8' | -7.67948 | 1.14E-24 |
| G93A_EOM_D/G93A_Dia_D | 22441 | 'Xlr' | 8.194757 | 3.42E-14 | 15402 | 'Hoxa5' | -7.48247 | 7.01E-97 |
| G93A_EOM_D/G93A_Dia_D | 18740 | 'Pitx1' | 8.055282 | 7.40E-32 | 56222 | 'Cited4' | -7.31288 | 4.35E-10 |
| G93A_EOM_D/G93A_Dia_D | 16371 | 'Irx1' | 7.888743 | 2.58E-24 | 15425 | 'Hoxc6' | -6.80959 | 1.96E-226 |
| G93A_EOM_D/G93A_Dia_D | 70691 | '3830403N18Rik' | 6.906891 | 3.65E-06 | 15404 | 'Hoxa7' | -6.71195 | 5.39E-58 |
| G93A_EOM_D/G93A_Dia_D | 56533 | 'Rgs17' | 6.78136 | 1.59E-45 | 15401 | 'Hoxa4' | -6.45943 | 7.59E-50 |
| G93A_EOM_D/G93A_Dia_D | 12159 | 'Bmp4' | 6.547996 | 1.59E-55 | 14537 | 'Gcnt1' | -6.42066 | 4.37E-57 |
| G93A_EOM_D/G93A_Dia_D | 22403 | 'Wisp2' | 5.8472 | 9.59E-197 | 21380 | 'Tbx1' | -6.33053 | 2.64E-155 |
| G93A_EOM_D/G93A_Dia_D | 242050 | 'Igsf10' | 5.807355 | 3.23E-47 | 15424 | 'Hoxc5' | -6.05528 | 1.26E-33 |
| G93A_EOM_D/G93A_Dia_D | 20319 | 'Sfrp2' | 5.709658 | 6.89E-26 | 15400 | 'Hoxa3' | -5.94953 | 2.04E-41 |
| G93A_EOM_D/G93A_Dia_D | 22355 | 'Vipr2' | 5.672425 | 1.79E-16 | 15399 | 'Hoxa2' | -5.93074 | 5.08E-15 |
| G93A_EOM_D/G93A_Dia_D | 208898 | 'Unc13c' | 5.61471 | 3.03E-39 | 545611 | 'Fam205a2' | -5.42626 | 5.31E-08 |
| G93A_EOM_D/G93A_Dia_D | 11695 | 'Alx4' | 5.613676 | 0 | 209448 | 'Hoxc10' | -5.39232 | 2.20E-04 |
| G93A_EOM_D/G93A_Dia_D | 20257 | 'Stmn2' | 5.595851 | 3.41E-88 | 16814 | 'Lbx1' | -5.33681 | 3.66E-213 |
| G93A_EOM_D/G93A_Dia_D | 11535 | 'Adm' | 5.577429 | 1.05E-11 | 15405 | 'Hoxa9' | -5.16993 | 3.55E-06 |
| G93A_EOM_D/G93A_Dia_D | 217843 | 'Unc79' | 5.523562 | 1.89E-18 | 13628 | 'Eef1a2' | -5.02975 | 3.73E-09 |
| G93A_EOM_D/G93A_Dia_D | 12023 | 'Barx2' | 5.485904 | 2.32E-93 | 140781 | 'Myh7' | -5.00355 | 0 |
| G93A_EOM_D/G93A_Dia_D | 20377 | 'Sfrp1' | 5.426265 | 1.16E-08 | 15394 | 'Hoxa1' | -4.73335 | 6.79E-26 |
| G93A_EOM_D/G93A_Dia_D | 14049 | 'Eya2' | 5.423466 | 5.40E-111 | 241431 | 'Xirp2' | -4.59694 | 0 |
