## Supplementary material for "Distinct transcriptomic profile of satellite cells contributes to preservation of neuromuscular junctions in extraocular muscles of ALS mice": Figure 6-Source Data 2

**Figure 6-Source Data 2. Top 20 differentially expressed genes comparing G93A to WT SCs of the same muscle origin cultured in growth and differentiation medium.** Log2FC and FDR are shown and genes are ranked according to log2FC. Column 2-5 are genes expressed higher in G93A SCs and Column 6-9 are genes expressed lower in G93A SCs.

| Group of comparison | Gene ID (higher) | Gene Symbol | log2 | FDR | Gene ID (lower) | Gene Symbol | log2 | FDR |
| --- | --- | --- | --- | --- | --- | --- | --- | --- |
| G93A_HL_G/WT_HL_G | 105246618 | 'Gm41885' | 11.10787 | 0 | 434233 | 'Ppp1ccb' | -9.02514 | 1.78E-37 |
| G93A_HL_G/WT_HL_G | 105247050 | 'Gm42226' | 10.82893 | 3.98E-196 | 1E+08 | 'Gm10045' | -8.04939 | 2.55E-131 |
| G93A_HL_G/WT_HL_G | 108169061 | 'Gm46918' | 9.548822 | 2.20E-33 | 621832 | 'Nutf2-ps2' | -7.88264 | 4.30E-07 |
| G93A_HL_G/WT_HL_G | 101056102 | 'Gm29779' | 9.408967 | 0 | 76681 | 'Trim12a' | -6.82018 | 1.67E-08 |
| G93A_HL_G/WT_HL_G | 319192 | 'Hist2h2aa2' | 8.219169 | 1.21E-07 | 545490 | 'Zfp973' | -6.71425 | 1.56E-06 |
| G93A_HL_G/WT_HL_G | 108167806 | 'Gm46221' | 8.159871 | 2.85E-31 | 1.03E+08 | 'LOC102639653' | -6.56986 | 4.05E-19 |
| G93A_HL_G/WT_HL_G | 105244980 | 'Gm40498' | 7.83041 | 0 | 170757 | 'Adgrl4' | -6.55459 | 4.05E-19 |
| G93A_HL_G/WT_HL_G | 105246904 | 'Gm42102' | 7.73471 | 3.14E-25 | 20957 | 'Sycp1' | -6.33985 | 1.21E-13 |
| G93A_HL_G/WT_HL_G | 278672 | 'Duxbl1' | 7.546894 | 1.03E-20 | 1E+08 | 'Gm14308' | -6.30378 | 1.14E-17 |
| G93A_HL_G/WT_HL_G | 111241 | 'Hmga1b' | 6.878562 | 3.08E-209 | 246256 | 'Fcgr4' | -6.26679 | 1.32E-04 |
| G93A_HL_G/WT_HL_G | 100039175 | 'Gm9780' | 6.870365 | 2.46E-05 | 107477 | 'Guca1b' | -5.93074 | 7.03E-05 |
| G93A_HL_G/WT_HL_G | 78376 | 'Sapcd1' | 6.72792 | 1.73E-04 | 1.08E+08 | 'Gm46386' | -4.83289 | 6.93E-08 |
| G93A_HL_G/WT_HL_G | 12353 | 'Car6' | 6.554589 | 3.39E-06 | 19944 | 'Rpl29' | -4.79977 | 0 |
| G93A_HL_G/WT_HL_G | 105244006 | 'Gm39701' | 6.388201 | 0 | 17921 | 'Myo7a' | -4.69515 | 1.06E-162 |
| G93A_HL_G/WT_HL_G | 547349 | 'LOC547349' | 6.285402 | 9.00E-05 | 1.08E+08 | 'LOC108168962' | -4.58496 | 8.22E-07 |
| G93A_HL_G/WT_HL_G | 109697 | 'Cpa1' | 5.954196 | 9.00E-05 | 67425 | 'Eps8l1' | -4.5025 | 4.92E-10 |
| G93A_HL_G/WT_HL_G | 105245547 | 'Gm40991' | 5.756668 | 0 | 228846 | 'D630003M21Rik' | -4.24793 | 8.50E-04 |
| G93A_HL_G/WT_HL_G | 665902 | 'Zscan4f' | 5.643856 | 1.73E-04 | 56642 | 'Ankrd2' | -4.18227 | 0 |
| G93A_HL_G/WT_HL_G | 105244343 | 'Gm39972' | 5.522528 | 0 | 19378 | 'Aldh1a2' | -4.16993 | 9.29E-08 |
| G93A_HL_G/WT_HL_G | 105246807 | 'Gm42031' | 5.217231 | 1.71E-25 | 69700 | 'Col22a1' | -4.08746 | 3.75E-05 |
| G93A_Dia_G/WT_Dia_G | 433182 | 'Eno1b' | 9.524103 | 0 | 11615 | 'Gm4737' | -12.3886 | 0 |
| G93A_Dia_G/WT_Dia_G | 105247050 | 'Gm42226' | 9.517669 | 9.85E-79 | 111241 | 'Hmga1b' | -11.5665 | 2.85E-240 |
| G93A_Dia_G/WT_Dia_G | 101056102 | 'Gm29779' | 9.048033 | 0 | 1E+08 | 'Gm2427' | -8.11374 | 2.05E-19 |
| G93A_Dia_G/WT_Dia_G | 621832 | 'Nutf2-ps2' | 7.139551 | 2.14E-04 | 545490 | 'Zfp973' | -7.46761 | 3.17E-11 |
| G93A_Dia_G/WT_Dia_G | 112422 | 'Zfp979' | 6.584963 | 1.52E-11 | 319192 | 'Hist2h2aa2' | -7.16993 | 7.74E-04 |
| G93A_Dia_G/WT_Dia_G | 50530 | 'Mfap5' | 6.266787 | 2.14E-04 | 1E+08 | 'Tmem254c' | -6.91886 | 3.42E-07 |
| G93A_Dia_G/WT_Dia_G | 71111 | 'Gpr39' | 4.906891 | 7.56E-04 | 668039 | 'Gm14434' | -6.85798 | 1.79E-26 |
| G93A_Dia_G/WT_Dia_G | 13179 | 'Dcn' | 4.459432 | 4.65E-05 | 20957 | 'Sycp1' | -6.58496 | 1.79E-16 |
| G93A_Dia_G/WT_Dia_G | 101488212 | 'Evi2' | 4.392317 | 4.05E-04 | 50874 | 'Tmod4' | -6.36632 | 9.26E-13 |
| G93A_Dia_G/WT_Dia_G | 21892 | 'Tll1' | 4.392317 | 2.74E-04 | 1.07E+08 | 'Gm45935' | -5.16993 | 6.64E-07 |
| G93A_Dia_G/WT_Dia_G | 18383 | 'Tnfrsf11b' | 4.129283 | 4.95E-04 | 18106 | 'Cd244a' | -5.16993 | 5.91E-10 |
| G93A_Dia_G/WT_Dia_G | 434215 | 'Lrrc32' | 3.824428 | 1.94E-24 | 57738 | 'Slc15a2' | -4.75489 | 9.04E-04 |
| G93A_Dia_G/WT_Dia_G | 105245604 | 'Gm41035' | 3.817832 | 1.50E-54 | 216984 | 'Evi2b' | -4.39232 | 7.73E-04 |
| G93A_Dia_G/WT_Dia_G | 12424 | 'Cck' | 3.509555 | 1.42E-04 | 74438 | 'Clvs1' | -3.70044 | 4.20E-07 |
| G93A_Dia_G/WT_Dia_G | 242384 | 'Lingo2' | 3.5025 | 3.94E-04 | 17906 | 'Myl2' | -3.52301 | 1.35E-25 |
| G93A_Dia_G/WT_Dia_G | 105245011 | 'Gm40525' | 3.415037 | 4.33E-04 | 270192 | 'Rab6b' | -3.51457 | 1.44E-12 |
| G93A_Dia_G/WT_Dia_G | 12983 | 'Csf2rb' | 3.415037 | 7.47E-04 | 170745 | 'Xpnpep2' | -3.39232 | 2.69E-05 |
| G93A_Dia_G/WT_Dia_G | 732521 | 'Gt(pU21)140Imeg' | 3.247928 | 2.86E-08 | 20465 | 'Sim2' | -3.3505 | 7.41E-06 |
| G93A_Dia_G/WT_Dia_G | 244416 | 'Ppp1r3b' | 2.969626 | 1.37E-12 | 59011 | 'Myoz1' | -3.21501 | 2.65E-10 |
| G93A_Dia_G/WT_Dia_G | 105244980 | 'Gm40498' | 2.92327 | 7.59E-110 | 71841 | 'Tcp11x2' | -3.05311 | 7.92E-09 |
| G93A_EOM_G/WT_EOM_G | 319192 | 'Hist2h2aa2' | 7.199672 | 7.10E-04 | 18811 | 'Prl2c2' | -7.45943 | 2.29E-06 |
| G93A_EOM_G/WT_EOM_G | 100040944 | 'Gm3055' | 5.459432 | 7.09E-04 | 18812 | 'Prl2c3' | -7.2854 | 8.33E-06 |
| G93A_EOM_G/WT_EOM_G | 668039 | 'Gm14434' | 5.087463 | 6.86E-14 | 504193 | 'Npcd' | -6.80735 | 1.28E-27 |
| G93A_EOM_G/WT_EOM_G | 101488212 | 'Evi2' | 4.392317 | 3.79E-04 | 619441 | 'Tnfsfm13' | -5.2854 | 3.83E-04 |
| G93A_EOM_G/WT_EOM_G | 100043876 | 'Gm4705' | 3.465381 | 8.29E-04 | 13592 | 'Ebf2' | -4.75489 | 7.97E-05 |
| G93A_EOM_G/WT_EOM_G | 56216 | 'Stx1b' | 2.954196 | 2.63E-04 | 216984 | 'Evi2b' | -4.52356 | 3.83E-04 |
| G93A_EOM_G/WT_EOM_G | 60596 | 'Gucy1a1' | 2.681824 | 2.20E-09 | 14537 | 'Gcnt1' | -4.45943 | 5.70E-05 |
| G93A_EOM_G/WT_EOM_G | 83558 | 'Tex11' | 2.61891 | 6.10E-04 | 1.01E+08 | 'LOC100862455' | -3.92348 | 0 |
| G93A_EOM_G/WT_EOM_G | 14264 | 'Fmod' | 2.590622 | 3.63E-90 | 67425 | 'Eps8l1' | -3.77259 | 7.25E-07 |
| G93A_EOM_G/WT_EOM_G | 333182 | 'Cox6b2' | 2.556634 | 1.51E-04 | 14563 | 'Gdf5' | -3.40599 | 2.35E-04 |
| G93A_EOM_G/WT_EOM_G | 69623 | 'Zfp33b' | 2.294183 | 3.67E-08 | 1E+08 | 'Gm14308' | -3.27716 | 2.11E-50 |
| G93A_EOM_G/WT_EOM_G | 108168045 | 'Gm46386' | 1.983512 | 1.02E-05 | 226691 | 'Ifi207' | -3 | 5.48E-06 |
| G93A_EOM_G/WT_EOM_G | 12291 | 'Cacna1g' | 1.954196 | 2.67E-05 | 241431 | 'Xirp2' | -2.78241 | 4.37E-112 |
| G93A_EOM_G/WT_EOM_G | 14747 | 'Cmklr1' | 1.934577 | 2.17E-33 | 14858 | 'Gsta2' | -2.68557 | 2.88E-16 |
| G93A_EOM_G/WT_EOM_G | 22403 | 'Wisp2' | 1.845769 | 1.88E-18 | 17751 | 'Mt3' | -2.63452 | 5.58E-05 |
| G93A_EOM_G/WT_EOM_G | 12159 | 'Bmp4' | 1.801824 | 3.21E-19 | 1.03E+08 | 'Gm21992' | -2.5502 | 1.40E-06 |
| G93A_EOM_G/WT_EOM_G | 278672 | 'Duxbl1' | 1.791892 | 2.31E-06 | 110893 | 'Slc8a3' | -2.38702 | 1.61E-07 |
| G93A_EOM_G/WT_EOM_G | 213436 | 'Rtl3' | 1.778306 | 6.89E-27 | 75668 | 'Rasl10a' | -2.35252 | 6.70E-04 |
| G93A_EOM_G/WT_EOM_G | 17242 | 'Mdk' | 1.768536 | 6.59E-05 | 12372 | 'Casq1' | -2.34915 | 8.97E-104 |
| G93A_EOM_G/WT_EOM_G | 208117 | 'Aph1b' | 1.762961 | 9.62E-17 | 1.03E+08 | 'LOC102633156' | -2.22517 | 4.24E-15 |
| G93A_HL_D/WT_HL_D | 15077 | 'Hist2h3c1' | 10.65284 | 3.78E-46 | 1E+08 | 'Gm14305' | -7.47573 | 9.83E-13 |
| G93A_HL_D/WT_HL_D | 105247050 | 'Gm42226' | 10.36632 | 7.25E-135 | 20957 | 'Sycp1' | -6.14975 | 2.63E-11 |
| G93A_HL_D/WT_HL_D | 621832 | 'Nutf2-ps2' | 9.079485 | 8.92E-16 | 107477 | 'Guca1b' | -6.10852 | 4.25E-05 |
| G93A_HL_D/WT_HL_D | 101056102 | 'Gm29779' | 9.013497 | 0 | 1.08E+08 | 'Gm46386' | -6 | 1.37E-09 |
| G93A_HL_D/WT_HL_D | 105246807 | 'Gm42031' | 8.900867 | 5.47E-35 | 1.01E+08 | 'Tpbgl' | -5.93074 | 1.88E-08 |
| G93A_HL_D/WT_HL_D | 105244980 | 'Gm40498' | 8.628027 | 0 | 545649 | 'Gm13276' | -5.90689 | 9.51E-04 |
| G93A_HL_D/WT_HL_D | 14559 | 'Gdf1' | 8.459432 | 2.39E-22 | 19944 | 'Rpl29' | -5.72484 | 0 |
| G93A_HL_D/WT_HL_D | 102640717 | 'Gm36718' | 8.005625 | 2.34E-11 | 76681 | 'Trim12a' | -5.58496 | 9.51E-04 |
| G93A_HL_D/WT_HL_D | 12353 | 'Car6' | 7.906891 | 6.90E-15 | 22064 | 'Trpc2' | -5.52356 | 4.80E-07 |
| G93A_HL_D/WT_HL_D | 105244006 | 'Gm39701' | 7.65422 | 0 | 1.08E+08 | 'Gm46290' | -5.49185 | 7.91E-05 |
| G93A_HL_D/WT_HL_D | 111241 | 'Hmga1b' | 7.584963 | 4.34E-167 | 1.03E+08 | 'Gm32687' | -5.3339 | 2.98E-12 |
| G93A_HL_D/WT_HL_D | 105246618 | 'Gm41885' | 7.426265 | 0 | 1E+08 | 'Gm10045' | -5.30223 | 4.72E-80 |
| G93A_HL_D/WT_HL_D | 108167809 | 'Gm46223' | 7.285402 | 6.91E-15 | 17933 | 'Myt1l' | -5 | 6.32E-09 |
| G93A_HL_D/WT_HL_D | 20389 | 'Sftpc' | 7.066089 | 1.08E-04 | 17921 | 'Myo7a' | -4.71049 | 1.45E-81 |
| G93A_HL_D/WT_HL_D | 108167806 | 'Gm46221' | 6.894818 | 6.08E-12 | 545490 | 'Zfp973' | -4.63421 | 1.44E-07 |
| G93A_HL_D/WT_HL_D | 105245547 | 'Gm40991' | 6.704223 | 0 | 668039 | 'Gm14434' | -4.52356 | 7.90E-05 |
| G93A_HL_D/WT_HL_D | 14167 | 'Fgf12' | 6.087463 | 1.69E-13 | 1.08E+08 | 'LOC108168962' | -4.49185 | 5.70E-06 |
| G93A_HL_D/WT_HL_D | 108168162 | 'Gm43305' | 6.008777 | 0 | 21892 | 'Tll1' | -4.45943 | 4.25E-05 |
| G93A_HL_D/WT_HL_D | 105246138 | 'Gm41476' | 5.918863 | 1.44E-27 | 434215 | 'Lrrc32' | -4.45346 | 4.88E-35 |
| G93A_HL_D/WT_HL_D | 105244343 | 'Gm39972' | 5.846753 | 0 | 16516 | 'Kcnj15' | -4.39232 | 3.58E-04 |
| G93A_Dia_D/WT_Dia_D | 101056102 | 'Gm29779' | 8.659401 | 0 | 11615 | 'Gm4737' | -11.4767 | 1.29E-291 |
| G93A_Dia_D/WT_Dia_D | 13999 | 'Gm14288' | 8.252665 | 3.50E-05 | 15077 | 'Hist2h3c1' | -11.1805 | 7.13E-68 |
| G93A_Dia_D/WT_Dia_D | 105247050 | 'Gm42226' | 7.906891 | 2.42E-24 | 111241 | 'Hmga1b' | -10.3095 | 2.10E-95 |
| G93A_Dia_D/WT_Dia_D | 433182 | 'Eno1b' | 7.389327 | 0 | 1E+08 | 'Gm2427' | -7.66534 | 1.62E-13 |
| G93A_Dia_D/WT_Dia_D | 12797 | 'Cnn1' | 6.149747 | 1.26E-04 | 319189 | 'Hist2h2bb' | -7.63662 | 4.36E-04 |
| G93A_Dia_D/WT_Dia_D | 20706 | 'Serpinb9b' | 6.066089 | 5.13E-06 | 1E+08 | 'Gm14308' | -6.97728 | 1.22E-27 |
| G93A_Dia_D/WT_Dia_D | 668039 | 'Gm14434' | 5.554589 | 5.15E-10 | 1.08E+08 | 'Gm46221' | -6.76818 | 3.42E-11 |
| G93A_Dia_D/WT_Dia_D | 22064 | 'Trpc2' | 5.491853 | 3.79E-07 | 1E+08 | 'Gm9780' | -6.44294 | 2.32E-04 |
| G93A_Dia_D/WT_Dia_D | 71111 | 'Gpr39' | 5.247928 | 6.67E-05 | 20957 | 'Sycp1' | -6.22882 | 2.39E-12 |
| G93A_Dia_D/WT_Dia_D | 50530 | 'Mfap5' | 4.964501 | 1.51E-13 | 667780 | 'Gm13871' | -6.08746 | 4.36E-04 |
| G93A_Dia_D/WT_Dia_D | 100043034 | 'Rex2' | 4.954196 | 8.35E-04 | 320825 | 'Samd5' | -5 | 8.20E-04 |
| G93A_Dia_D/WT_Dia_D | 16775 | 'Lama4' | 4.643856 | 3.79E-07 | 56533 | 'Rgs17' | -4.75489 | 1.30E-10 |
| G93A_Dia_D/WT_Dia_D | 434215 | 'Lrrc32' | 3.862496 | 1.42E-44 | 14559 | 'Gdf1' | -4.5253 | 2.99E-83 |
| G93A_Dia_D/WT_Dia_D | 21892 | 'Tll1' | 3.61471 | 4.01E-09 | 57738 | 'Slc15a2' | -4.35755 | 2.76E-10 |
| G93A_Dia_D/WT_Dia_D | 105245604 | 'Gm41035' | 3.581795 | 4.08E-33 | 12350 | 'Car3' | -4.12722 | 5.46E-38 |
| G93A_Dia_D/WT_Dia_D | 18383 | 'Tnfrsf11b' | 3.491853 | 2.76E-04 | 270192 | 'Rab6b' | -3.12199 | 8.29E-21 |
| G93A_Dia_D/WT_Dia_D | 12819 | 'Col15a1' | 3.380822 | 7.31E-20 | 278672 | 'Duxbl1' | -2.9542 | 2.46E-14 |
| G93A_Dia_D/WT_Dia_D | 26564 | 'Ror2' | 3.273018 | 8.32E-04 | 20660 | 'Sorl1' | -2.48543 | 8.33E-11 |
| G93A_Dia_D/WT_Dia_D | 12424 | 'Cck' | 3.218424 | 3.97E-07 | 1.01E+08 | 'Fam205a3' | -2.42626 | 4.88E-04 |
| G93A_Dia_D/WT_Dia_D | 105246904 | 'Gm42102' | 3.105534 | 1.30E-15 | 380713 | 'Scarf1' | -2.36678 | 1.77E-06 |
| G93A_EOM_D/WT_EOM_D | 15077 | 'Hist2h3c1' | 10.1472 | 1.87E-34 | 14559 | 'Gdf1' | -7.2384 | 6.17E-10 |
| G93A_EOM_D/WT_EOM_D | 100328588 | 'Il4i1b' | 5.754888 | 3.99E-06 | 504193 | 'Npcd' | -6.20945 | 5.54E-18 |
| G93A_EOM_D/WT_EOM_D | 625591 | 'Cldn34c2' | 3.788496 | 1.35E-04 | 1.08E+08 | 'Gm46290' | -6.06609 | 1.14E-07 |
| G93A_EOM_D/WT_EOM_D | 100043876 | 'Gm4705' | 3.079727 | 3.92E-05 | 14204 | 'Il4i1' | -5.90689 | 5.36E-06 |
| G93A_EOM_D/WT_EOM_D | 668039 | 'Gm14434' | 2.392317 | 4.44E-07 | 1.01E+08 | 'Fam205a3' | -5.45943 | 1.63E-08 |
| G93A_EOM_D/WT_EOM_D | 67464 | 'Entpd4' | 2.2892 | 1.14E-34 | 97114 | 'Hist2h3c2' | -4.72204 | 7.91E-33 |
| G93A_EOM_D/WT_EOM_D | 54123 | 'Irf7' | 2.219313 | 1.36E-11 | 16185 | 'Il2rb' | -4.52356 | 1.28E-04 |
| G93A_EOM_D/WT_EOM_D | 24110 | 'Usp18' | 2.169925 | 9.97E-04 | 1.01E+08 | 'Zfp965' | -3.7761 | 3.01E-06 |
| G93A_EOM_D/WT_EOM_D | 107303348 | 'Gm45935' | 2.159199 | 8.78E-06 | 67425 | 'Eps8l1' | -3.70044 | 1.28E-10 |
| G93A_EOM_D/WT_EOM_D | 66443 | 'Tnfaip8l1' | 2.036837 | 1.33E-05 | 1.03E+08 | 'Gm14296' | -3.0902 | 7.74E-31 |
| G93A_EOM_D/WT_EOM_D | 235130 | 'Adamts15' | 2.016302 | 7.92E-12 | 74318 | 'Hopx' | -2.84398 | 2.10E-42 |
| G93A_EOM_D/WT_EOM_D | 237387 | 'Lrrc3' | 1.906891 | 8.63E-04 | 1E+08 | 'Gm14305' | -2.64909 | 2.71E-11 |
| G93A_EOM_D/WT_EOM_D | 100042807 | 'Eif3j2' | 1.882086 | 1.59E-116 | 21897 | 'Tlr1' | -2.39941 | 4.59E-10 |
| G93A_EOM_D/WT_EOM_D | 72962 | 'Tymp' | 1.881356 | 3.08E-05 | 667962 | 'Zfp966' | -2.383 | 1.38E-18 |
| G93A_EOM_D/WT_EOM_D | 69623 | 'Zfp33b' | 1.880418 | 1.98E-05 | 240025 | 'Dact2' | -2.34792 | 5.05E-04 |
| G93A_EOM_D/WT_EOM_D | 241688 | 'Dzank1' | 1.874469 | 1.41E-04 | 380728 | 'Kcnh4' | -2.28951 | 5.32E-04 |
| G93A_EOM_D/WT_EOM_D | 434233 | 'Ppp1ccb' | 1.873432 | 1.33E-22 | 1.03E+08 | 'LOC102638183' | -2.12853 | 8.45E-04 |
| G93A_EOM_D/WT_EOM_D | 100504263 | '2210418O10Rik' | 1.813988 | 3.35E-07 | 12984 | 'Csf2rb2' | -2.10434 | 5.33E-04 |
| G93A_EOM_D/WT_EOM_D | 100044627 | 'LOC100044627' | 1.79773 | 3.62E-119 | 71690 | 'Esm1' | -1.94753 | 2.09E-04 |
| G93A_EOM_D/WT_EOM_D | 100038513 | 'Gm14149' | 1.745954 | 9.44E-18 | 1E+08 | 'Gm3558' | -1.92811 | 2.19E-05 |
