## Supplementary material for "Distinct transcriptomic profile of satellite cells contributes to preservation of neuromuscular junctions in extraocular muscles of ALS mice": Figure 6-Source Data 3

**Figure 6-Source Data 3. Top 20 differentially expressed genes comparing G93A hindlimb and diaphragm SCs with 3-day NaBu treatment to those without.** Log2FC and FDR are shown and genes are ranked according to log2FC. Column 2-5 are genes expressed higher in NaBu treated group and Column 6-9 are genes expressed lower in NaBu treated group.

| Group of comparison | Gene ID (up) | Gene Symbol | log2 | FDR | Gene ID (down) | Gene Symbol | log2 | FDR |
| --- | --- | --- | --- | --- | --- | --- | --- | --- |
| G93A_HL_D_NaBu3/G93A_HL_D | 545490 | 'Zfp973' | 7.954196 | 7.26E-17 | 100503949 | 'Zfp965' | -7.77479 | 1.29E-14 |
| G93A_HL_D_NaBu3/G93A_HL_D | 14999 | 'H2-DMb1' | 7.622052 | 7.75E-13 | 319192 | 'Hist2h2aa2' | -7.09803 | 3.02E-04 |
| G93A_HL_D_NaBu3/G93A_HL_D | 668039 | 'Gm14434' | 6.523562 | 2.40E-22 | 13386 | 'Dlk1' | -6.82018 | 3.95E-10 |
| G93A_HL_D_NaBu3/G93A_HL_D | 54120 | 'Gipc2' | 6.108524 | 4.50E-04 | 100040766 | 'Mroh2a' | -5.58496 | 7.82E-10 |
| G93A_HL_D_NaBu3/G93A_HL_D | 102294 | 'Cyp4v3' | 5.954196 | 1.07E-09 | 15221 | 'Foxd3' | -5.20945 | 2.41E-08 |
| G93A_HL_D_NaBu3/G93A_HL_D | 110095 | 'Pygl' | 5.930737 | 7.95E-17 | 383563 | 'Gpr25' | -5.12928 | 5.85E-04 |
| G93A_HL_D_NaBu3/G93A_HL_D | 20981 | 'Syt3' | 5.84549 | 1.07E-15 | 14611 | 'Gja3' | -4.97728 | 1.44E-15 |
| G93A_HL_D_NaBu3/G93A_HL_D | 20732 | 'Spint1' | 5.807355 | 7.31E-07 | 320825 | 'Samd5' | -4.93074 | 1.71E-07 |
| G93A_HL_D_NaBu3/G93A_HL_D | 74134 | 'Cyp2s1' | 5.523562 | 1.39E-06 | 242022 | 'Frem2' | -4.79307 | 0 |
| G93A_HL_D_NaBu3/G93A_HL_D | 623230 | 'Tmem200b' | 5.357552 | 3.49E-05 | 665596 | 'Hist1h2bq' | -4.7907 | 7.47E-15 |
| G93A_HL_D_NaBu3/G93A_HL_D | 242700 | 'Ifnlr1' | 5.285402 | 1.47E-08 | 631797 | 'Fer1l6' | -4.58496 | 2.25E-06 |
| G93A_HL_D_NaBu3/G93A_HL_D | 269295 | 'Rtn4rl2' | 5.239466 | 2.37E-28 | 668303 | 'Kif26a' | -4.45396 | 2.01E-78 |
| G93A_HL_D_NaBu3/G93A_HL_D | 20713 | 'Serpini1' | 5.169925 | 1.10E-10 | 244698 | 'Hephl1' | -4.42321 | 8.59E-105 |
| G93A_HL_D_NaBu3/G93A_HL_D | 14961 | 'H2-Ab1' | 5.161888 | 5.92E-19 | 73748 | 'Gadl1' | -4.40939 | 1.33E-12 |
| G93A_HL_D_NaBu3/G93A_HL_D | 15377 | 'Foxa3' | 5.129283 | 4.50E-04 | 22411 | 'Wnt11' | -4.39232 | 4.32E-18 |
| G93A_HL_D_NaBu3/G93A_HL_D | 69325 | '1700012B09Rik' | 5.08274 | 4.89E-09 | 13426 | 'Dync1i1' | -4.32992 | 6.60E-21 |
| G93A_HL_D_NaBu3/G93A_HL_D | 277328 | 'Trpa1' | 5 | 2.45E-06 | 11448 | 'Chrne' | -4.31159 | 1.26E-09 |
| G93A_HL_D_NaBu3/G93A_HL_D | 108167568 | 'Gm46063' | 4.97728 | 4.89E-09 | 93960 | 'Nkd1' | -4.23095 | 2.15E-30 |
| G93A_HL_D_NaBu3/G93A_HL_D | 380863 | 'Tmem171' | 4.93546 | 3.93E-10 | 69698 | 'Slc52a3' | -4.22239 | 6.25E-07 |
| G93A_HL_D_NaBu3/G93A_HL_D | 12475 | 'Cd14' | 4.874469 | 1.33E-06 | 245026 | 'Col6a6' | -4.1964 | 2.66E-15 |
| G93A_Dia_D_NaBu3/G93A_Dia_D | 68458 | 'Ppp1r14a' | 8.724514 | 1.27E-10 | 100503949 | 'Zfp965' | -7.9542 | 7.50E-17 |
| G93A_Dia_D_NaBu3/G93A_Dia_D | 545490 | 'Zfp973' | 7.774787 | 7.92E-15 | 27528 | 'Nrep' | -7.41785 | 6.14E-20 |
| G93A_Dia_D_NaBu3/G93A_Dia_D | 668039 | 'Gm14434' | 6.569856 | 1.00E-22 | 504193 | 'Npcd' | -7.36632 | 6.75E-44 |
| G93A_Dia_D_NaBu3/G93A_Dia_D | 140743 | 'Rem2' | 6.066089 | 9.80E-07 | 100039830 | 'Gm2446' | -7.2854 | 2.32E-07 |
| G93A_Dia_D_NaBu3/G93A_Dia_D | 103098 | 'Slc6a15' | 5.894818 | 2.58E-20 | 237759 | 'Col23a1' | -7.18322 | 1.00E-138 |
| G93A_Dia_D_NaBu3/G93A_Dia_D | 14999 | 'H2-DMb1' | 5.270529 | 2.92E-11 | 319192 | 'Hist2h2aa2' | -7.10852 | 4.57E-04 |
| G93A_Dia_D_NaBu3/G93A_Dia_D | 103140 | 'Gstt3' | 5.209453 | 8.90E-19 | 13386 | 'Dlk1' | -6.47573 | 7.07E-09 |
| G93A_Dia_D_NaBu3/G93A_Dia_D | 14961 | 'H2-Ab1' | 5.116864 | 3.12E-18 | 73748 | 'Gadl1' | -6.40939 | 5.26E-15 |
| G93A_Dia_D_NaBu3/G93A_Dia_D | 269295 | 'Rtn4rl2' | 5.087463 | 1.09E-17 | 11514 | 'Adcy8' | -6.10852 | 5.16E-19 |
| G93A_Dia_D_NaBu3/G93A_Dia_D | 623230 | 'Tmem200b' | 4.807355 | 9.80E-04 | 383563 | 'Gpr25' | -6.08746 | 2.32E-07 |
| G93A_Dia_D_NaBu3/G93A_Dia_D | 242700 | 'Ifnlr1' | 4.807355 | 3.49E-06 | 11448 | 'Chrne' | -5.72792 | 5.88E-05 |
| G93A_Dia_D_NaBu3/G93A_Dia_D | 20732 | 'Spint1' | 4.790077 | 7.75E-09 | 22229 | 'Ucp3' | -5.70044 | 4.66E-07 |
| G93A_Dia_D_NaBu3/G93A_Dia_D | 50781 | 'Dkk3' | 4.754888 | 1.43E-08 | 242022 | 'Frem2' | -5.47478 | 0 |
| G93A_Dia_D_NaBu3/G93A_Dia_D | 14871 | 'Gstt1' | 4.657371 | 2.79E-19 | 142687 | 'Asb14' | -5.36923 | 3.08E-11 |
| G93A_Dia_D_NaBu3/G93A_Dia_D | 12495 | 'Entpd1' | 4.643856 | 6.56E-06 | 100040766 | 'Mroh2a' | -5.20945 | 7.94E-18 |
| G93A_Dia_D_NaBu3/G93A_Dia_D | 108115 | 'Slco4a1' | 4.596935 | 1.43E-44 | 433940 | 'Fam222a' | -5.08746 | 2.95E-05 |
| G93A_Dia_D_NaBu3/G93A_Dia_D | 232371 | 'C1rl' | 4.523562 | 5.28E-04 | 77018 | 'Col25a1' | -5.07631 | 0 |
| G93A_Dia_D_NaBu3/G93A_Dia_D | 18751 | 'Prkcb' | 4.498251 | 2.23E-42 | 320825 | 'Samd5' | -5 | 1.17E-04 |
| G93A_Dia_D_NaBu3/G93A_Dia_D | 50909 | 'C1ra' | 4.47032 | 1.45E-15 | 320292 | 'Rasgef1b' | -4.64386 | 2.31E-04 |
| G93A_Dia_D_NaBu3/G93A_Dia_D | 277328 | 'Trpa1' | 4.459432 | 2.31E-05 | 668303 | 'Kif26a' | -4.62727 | 2.96E-53 |
