## Supplementary material for "Distinct transcriptomic profile of satellite cells contributes to preservation of neuromuscular junctions in extraocular muscles of ALS mice": Figure 6-Source Data 4

**Figure 6-Source Data 4. Three subgroups of quiescent signature genes identified in the DEG lists comparing EOM SCs to diaphragm and hindlimb counterparts cultured in differentiation medium.** Log2FC and FDR are shown and genes are ranked according to relative abundance (TPM, from highest to lowest) in WT EOM SCs.

| Gene ID | Gene Symbol | log2 (G93A_EOM_D / G93A_Dia_D) | FDR (G93A_EOM_D / G93A_Dia_D) | log2 (G93A_EOM_D / G93A_HL_D) | FDR (G93A_EOM_D / G93A_HL_D) | log2 (WT_EOM_D / WT_Dia_D) | FDR (WT_EOM_D / WT_Dia_D) | log2 (WT_EOM_D / WT_HL_D) | FDR (WT_EOM_D / WT_HL_D) | Subgroup |
| --- | --- | --- | --- | --- | --- | --- | --- | --- | --- | --- |
| 17967 | 'Ncam1' | -0.722694752 | 0 | -0.950344543 | 0 | -0.495666614 | 0 | -0.95976572 | 0 | Most commonly used SC marker |
| 12555 | 'Cdh15' | 0.9140522 | 0 | 1.167660174 | 0 | 0.689232229 | 1.75E-214 | 1.048883223 | 0 | Most commonly used SC marker |
| 12389 | 'Cav1' | 1.883790494 | 0 | 3.028853114 | 0 | 2.398981153 | 0 | 2.62492624 | 0 | Most commonly used SC marker |
| 20971 | 'Sdc4' | 0.900610282 | 2.40E-51 | 0.468811835 | 1.16E-16 | 1.188927474 | 7.22E-111 | 1.333540728 | 2.45E-131 | Most commonly used SC marker |
| 14186 | 'Fgfr4' | 1.899200371 | 0 | 2.029829814 | 0 | 1.239660297 | 4.37E-137 | 1.766654634 | 3.66E-234 | Signaling pathways regulating pluripotency of stem cells (KEGG) |
| 56198 | 'Heyl' | 2.029747343 | 0 | 3.317490736 | 0 | 1.934313291 | 5.24E-278 | 2.903875558 | 0 | Notch signaling pathway (KEGG) |
| 18131 | 'Notch3' | 2.552069545 | 0 | 2.468498178 | 0 | 1.693148018 | 0 | 2.142132849 | 0 | Notch signaling pathway (KEGG) |
| 22329 | 'Vcam1' | 1.800081493 | 1.10E-139 | 2.065751647 | 4.54E-164 | 1.651363726 | 4.78E-108 | 2.063790012 | 9.28E-145 | Most commonly used SC marker |
| 17295 | 'Met' | 1.587369802 | 2.57E-259 | 1.634386428 | 1.29E-267 | 1.129678547 | 1.25E-118 | 1.047601033 | 4.85E-105 | Most commonly used SC marker |
| 20970 | 'Sdc3' | 1.359058803 | 1.20E-144 | 1.930929607 | 9.44E-235 | 1.118412508 | 1.39E-81 | 1.043971252 | 1.64E-70 | Most commonly used SC marker |
| 207521 | 'Dtx4' | 2.534663325 | 4.48E-294 | 3.225977807 | 0 | 2.249415595 | 1.89E-208 | 3.722168592 | 0 | Notch signaling pathway (KEGG) |
| 16600 | 'Klf4' | 1.526183089 | 1.67E-56 | 1.200967423 | 8.76E-39 | 1.180169033 | 1.43E-48 | 2.967108118 | 3.79E-162 | Signaling pathways regulating pluripotency of stem cells (KEGG) |
| 18509 | 'Pax7' | 1.75085112 | 2.29E-129 | 2.103093493 | 6.73E-172 | 1.301833768 | 2.44E-60 | 2.016500695 | 6.27E-121 | Most commonly used SC marker |
| 18129 | 'Notch2' | 1.071495591 | 3.51E-127 | 0.948201099 | 4.78E-102 | 0.438121112 | 8.96E-21 | 0.57810883 | 2.10E-33 | Notch signaling pathway (KEGG) |
| 12767 | 'Cxcr4' | 2.933288915 | 1.97E-80 | 1.988518436 | 4.10E-50 | 2.159446077 | 1.48E-41 | 1.497354102 | 2.08E-25 | Most commonly used SC marker |
| 18128 | 'Notch1' | 2.177538186 | 0 | 2.425465699 | 0 | 1.316332632 | 2.93E-109 | 1.824272712 | 1.18E-173 | Notch signaling pathway (KEGG) |
| 14366 | 'Fzd4' | 3.582579849 | 9.94E-150 | 3.011972642 | 1.28E-124 | 2.584962501 | 1.38E-85 | 2.738996129 | 2.77E-90 | Signaling pathways regulating pluripotency of stem cells (KEGG) |
| 18708 | 'Pik3r1' | 1.412843286 | 3.58E-65 | 1.430491152 | 2.58E-65 | 1.33219643 | 1.34E-55 | 1.173767068 | 5.28E-47 | Signaling pathways regulating pluripotency of stem cells (KEGG) |
| 12490 | 'Cd34' | 2.182864057 | 3.02E-35 | 2.521952703 | 2.81E-40 | 2.311300459 | 1.28E-29 | 3.379045066 | 3.26E-43 | Most commonly used SC marker |
| 12156 | 'Bmp2' | 2.807354922 | 1.21E-23 | 1.459431619 | 3.41E-10 | 2.996157934 | 9.03E-37 | 2.058893689 | 2.91E-25 | Signaling pathways regulating pluripotency of stem cells (KEGG) |
| 16000 | 'Igf1' | 1.621703048 | 1.30E-49 | 1.915988387 | 2.34E-61 | 1.483876376 | 2.89E-32 | 1.632268215 | 4.11E-36 | Signaling pathways regulating pluripotency of stem cells (KEGG) |
| 12159 | 'Bmp4' | 6.547996174 | 1.59E-55 | 9.355351096 | 1.10E-57 | 3.875780063 | 1.67E-23 | 5.060204634 | 2.78E-27 | Signaling pathways regulating pluripotency of stem cells (KEGG) |
| 16449 | 'Jag1' | 0.680931559 | 3.59E-06 | 2.360530934 | 4.28E-36 | 0.764187063 | 5.34E-06 | 2.126757142 | 5.03E-25 | Notch signaling pathway (KEGG) |
| 17681 | 'Msc' | 1.166249343 | 2.65E-05 | 4.029747343 | 9.78E-21 | 1.695145418 | 9.15E-06 | 3.502500341 | 6.39E-13 | Notch signaling pathway (KEGG) |
| 16880 | 'Lifr' | 1.573466862 | 5.78E-21 | 2.058893689 | 4.74E-29 | 0.535596423 | 8.08E-05 | 0.631355406 | 1.63E-07 | Signaling pathways regulating pluripotency of stem cells (KEGG) |
