## Supplementary material for "Distinct transcriptomic profile of satellite cells contributes to preservation of neuromuscular junctions in extraocular muscles of ALS mice": Figure 6-Source Data 5

**Figure 6-Source Data 5. Three subgroups of quiescent signature genes identified in the DEG lists comparing EOM SCs to diaphragm and hindlimb counterparts cultured in growth medium.** Log2FC and FDR are shown and genes are ranked according to relative abundance (TPM, from highest to lowest) in WT EOM SCs.

| Gene ID | Gene Symbol | log2 (G93A_EOM_G / G93A_Dia_G) | FDR (G93A_EOM_G / G93A_Dia_G) | log2 (G93A_EOM_G / G93A_HL_G) | FDR (G93A_EOM_G / G93A_HL_G) | log2 (WT_EOM_G / WT_Dia_G) | FDR (WT_EOM_G / WT_Dia_G) | log2 (WT_EOM_G / WT_HL_G) | FDR (WT_EOM_G / WT_HL_G) | Subgroup |
| --- | --- | --- | --- | --- | --- | --- | --- | --- | --- | --- |
| 16404 | 'Itga7' | -0.732040021 | 0 | -0.411778379 | 3.08E-107 | -0.757842542 | 0 | -0.888088934 | 0 | Most commonly used SC marker |
| 17967 | 'Ncam1' | -0.994515355 | 0 | -1.275303457 | 0 | -0.877788923 | 0 | -1.082663623 | 0 | Most commonly used SC marker |
| 17295 | 'Met' | 0.438041289 | 3.22E-54 | 0.691941157 | 6.49E-120 | 0.573943244 | 2.61E-103 | 0.457226899 | 8.56E-67 | Most commonly used SC marker |
| 14186 | 'Fgfr4' | -0.419752964 | 3.28E-34 | 0.61998138 | 1.22E-51 | -0.684823289 | 4.46E-61 | -0.688005538 | 3.90E-70 | Signaling pathways regulating pluripotency of stem cells (KEGG) |
| 22329 | 'Vcam1' | 0.61903987 | 7.58E-31 | 0.666648508 | 8.60E-35 | 0.76080163 | 3.19E-51 | 0.609059799 | 8.31E-35 | Most commonly used SC marker |
| 20971 | 'Sdc4' | 0.702526045 | 6.09E-20 | 0.862438161 | 7.06E-28 | 0.661026688 | 9.26E-29 | 0.812893479 | 5.78E-41 | Most commonly used SC marker |
| 18708 | 'Pik3r1' | 0.98622715 | 3.48E-37 | 0.788024195 | 1.66E-25 | 1.234169589 | 7.89E-55 | 1.00503559 | 2.60E-39 | Signaling pathways regulating pluripotency of stem cells (KEGG) |
| 56198 | 'Heyl' | -0.473792528 | 2.87E-23 | 0.793040814 | 3.93E-41 | -1.177538186 | 1.84E-58 | -0.68182404 | 2.83E-17 | Notch signaling pathway (KEGG) |
| 12490 | 'Cd34' | 1.164038158 | 1.19E-09 | 1.719099173 | 7.01E-17 | 2.007195501 | 4.50E-19 | 2.051138849 | 2.42E-20 | Most commonly used SC marker |
| 12159 | 'Bmp4' | 3.801823816 | 3.93E-46 | 3.234139307 | 2.12E-41 | 2.415037499 | 1.65E-10 | 2.415037499 | 4.46E-10 | Signaling pathways regulating pluripotency of stem cells (KEGG) |
| 16000 | 'Igf1' | 0.559427409 | 9.39E-05 | 0.830074999 | 2.75E-08 | 0.889296536 | 6.27E-07 | 0.648288436 | 2.01E-04 | Signaling pathways regulating pluripotency of stem cells (KEGG) |
