## Supplementary material for "Distinct transcriptomic profile of satellite cells contributes to preservation of neuromuscular junctions in extraocular muscles of ALS mice": Figure 6-Source Data 6

**Figure 6-Source Data 6.** **Three subgroups of quiescent signature genes identified in the DEG lists comparing G93A diaphragm and hindlimb SCs with and without 3-day NaBu treatment.** Log2FC and FDR are shown and genes are ranked according to relative abundance (TPM, from highest to lowest) in G93A HL SCs.

| Gene ID | Gene Symbol | log2  (G93A_Dia_D_NaBu3/ G93A_Dia_D) | FDR  (G93A_Dia_D_NaBu3/ G93A_Dia_D) | log2  (G93A_HL_D_NaBu3/ G93A_HL_D) | FDR  (G93A_HL_D_NaBu3/ G93A_HL_D) | Subgroup |
| --- | --- | --- | --- | --- | --- | --- |
| 16404 | 'Itga7' | 0.650830856 | 0 | 0.402116786 | 3.27E-239 | Most commonly used SC marker |
| 12558 | 'Cdh2' | 0.883481668 | 0 | 0.72382373 | 0 | Most commonly used SC marker |
| 19664 | 'Rbpj' | 1.78958022 | 0 | 1.370417941 | 6.59E-244 | Notch signaling pathway (KEGG) |
| 20971 | 'Sdc4' | 1.548482806 | 1.13E-100 | 1.273974872 | 1.46E-108 | Most commonly used SC marker |
| 12389 | 'Cav1' | 1.871324277 | 1.15E-179 | 1.474396499 | 2.94E-98 | Most commonly used SC marker |
| 20970 | 'Sdc3' | 1.708563097 | 5.04E-207 | 1.39093971 | 2.92E-127 | Most commonly used SC marker |
| 16600 | 'Klf4' | -0.447981598 | 1.10E-04 | -0.439764585 | 1.43E-04 | Signaling pathways regulating pluripotency of stem cells (KEGG) |
| 18131 | 'Notch3' | 1.29961699 | 8.41E-160 | 1.070668947 | 3.33E-94 | Notch signaling pathway (KEGG) |
| 17295 | 'Met' | 0.635061112 | 7.00E-19 | 0.815287782 | 1.87E-32 | Most commonly used SC marker |
| 18509 | 'Pax7' | 1.029559798 | 1.04E-21 | 1.057195402 | 4.12E-21 | Most commonly used SC marker |
| 22329 | 'Vcam1' | 2.138191231 | 5.49E-88 | 1.964566307 | 2.39E-58 | Most commonly used SC marker |
| 22417 | 'Wnt4' | -0.712901889 | 4.52E-04 | -0.939739475 | 9.95E-06 | Signaling pathways regulating pluripotency of stem cells (KEGG) |
| 12156 | 'Bmp2' | -1.662965013 | 2.86E-05 | -0.798366139 | 9.57E-04 | Signaling pathways regulating pluripotency of stem cells (KEGG) |
| 12490 | 'Cd34' | 3.35695548 | 1.93E-103 | 2.778973121 | 1.16E-53 | Most commonly used SC marker |
| 14366 | 'Fzd4' | 1.986060809 | 7.54E-08 | 1.807354922 | 1.24E-10 | Signaling pathways regulating pluripotency of stem cells (KEGG) |
| 12159 | 'Bmp4' | 2.349149564 | 3.92E-05 | 2.040414268 | 8.50E-07 | Signaling pathways regulating pluripotency of stem cells (KEGG) |
