## Supplementary material for "Distinct transcriptomic profile of satellite cells contributes to preservation of neuromuscular junctions in extraocular muscles of ALS mice": Figure 7-Source Data 1

**Figure 7-Source Data 1. Axon guidance related genes (KEGG) identified in EOM SC signature genes cultured in growth medium.** Log2FC and FDR are shown and genes are ranked according to relative abundance (TPM, from highest to lowest) in WT EOM SCs.

| Gene ID | Gene Symbol | log2 (G93A_EOM_G / G93A_Dia_G) | FDR (G93A_EOM_G / G93A_Dia_G) | log2 (G93A_EOM_G / G93A_HL_G) | FDR (G93A_EOM_G / G93A_HL_G) | log2 (WT_EOM_G / WT_Dia_G) | FDR (WT_EOM_G / WT_Dia_G) | log2 (WT_EOM_G / WT_HL_G) | FDR (WT_EOM_G / WT_HL_G) |
| --- | --- | --- | --- | --- | --- | --- | --- | --- | --- |
| 14677 | 'Gnai1' | 1.683757 | 0 | 2.368276 | 0 | 2.070479 | 0 | 1.702395 | 8.37E-298 |
| 17295 | 'Met' | 0.438041 | 3.22E-54 | 0.691941 | 6.49E-120 | 0.573943 | 2.61E-103 | 0.457227 | 8.56E-67 |
| 20315 | 'Cxcl12' | 1.153981 | 7.49E-76 | 3.673631 | 0 | 0.466182 | 1.69E-09 | 0.536571 | 3.34E-27 |
| 13642 | 'Efnb2' | 0.907784 | 8.03E-61 | 1.513553 | 3.35E-138 | 1.539351 | 9.08E-149 | 1.31719 | 2.47E-110 |
| 18803 | 'Plcg1' | 0.651426 | 3.24E-35 | 0.574743 | 3.89E-28 | 0.495665 | 1.65E-20 | 0.407058 | 1.47E-14 |
| 18186 | 'Nrp1' | 1.113468 | 1.10E-116 | 1.503415 | 1.10E-184 | 1.20926 | 2.66E-110 | 0.777519 | 4.07E-50 |
| 18845 | 'Plxna2' | 0.82834 | 2.99E-116 | 0.762803 | 9.00E-100 | 0.908909 | 9.84E-125 | 0.614381 | 2.92E-63 |
| 13846 | 'Ephb4' | 0.917538 | 7.09E-49 | 1.134056 | 3.30E-68 | 0.581322 | 7.79E-18 | 0.674249 | 4.85E-23 |
| 13835 | 'Epha1' | 3.611435 | 8.41E-172 | 2.6462 | 7.08E-127 | 2.756096 | 5.41E-131 | 1.37658 | 8.66E-52 |
| 18708 | 'Pik3r1' | 0.986227 | 3.48E-37 | 0.788024 | 1.66E-25 | 1.23417 | 7.89E-55 | 1.005036 | 2.60E-39 |
| 20352 | 'Sema4b' | 0.820043 | 1.31E-15 | 0.54374 | 3.23E-08 | 0.839806 | 1.15E-16 | 0.672265 | 1.23E-11 |
| 117606 | 'Boc' | 1.012175 | 3.33E-28 | 0.707466 | 1.10E-15 | 0.508237 | 5.55E-07 | 0.846785 | 4.83E-16 |
| 74769 | 'Pik3cb' | 1.027769 | 1.75E-19 | 0.524969 | 2.02E-07 | 1.217231 | 1.04E-28 | 1.075051 | 5.80E-25 |
| 18846 | 'Plxna3' | 0.784271 | 5.88E-23 | 1.012116 | 2.74E-36 | 0.56469 | 2.73E-11 | 0.622885 | 5.84E-11 |
| 12934 | 'Dpysl2' | 0.56718 | 5.61E-09 | 1.113595 | 1.60E-26 | 0.560146 | 2.43E-07 | 0.861256 | 8.47E-15 |
| 13848 | 'Ephb6' | 2.902703 | 2.98E-54 | 1.197446 | 7.13E-17 | 2.724893 | 1.48E-46 | 0.57289 | 3.44E-05 |
| 12167 | 'Bmpr1b' | 1.893913 | 6.31E-23 | 1.218535 | 8.92E-12 | 3.33787 | 6.24E-78 | 2.215013 | 5.83E-50 |
| 73181 | 'Nfatc4' | 2.971431 | 1.58E-82 | 4.785875 | 1.00E-121 | 2.459432 | 1.35E-29 | 3.704544 | 7.39E-45 |
| 65254 | 'Dpysl5' | 0.875194 | 3.42E-09 | 1.287577 | 1.52E-16 | 1.207488 | 1.08E-15 | 2.022649 | 3.42E-32 |
| 18707 | 'Pik3cd' | 0.576695 | 3.65E-04 | 0.929874 | 4.12E-08 | 0.776874 | 1.80E-06 | 1.112059 | 3.90E-11 |
| 13640 | 'Efna5' | 1.111283 | 6.64E-10 | 2.709921 | 7.03E-32 | 1.17433 | 2.35E-11 | 1.75269 | 3.82E-20 |
| 22418 | 'Wnt5a' | 2.604328 | 8.70E-28 | 1.623957 | 2.95E-15 | 3.493989 | 7.05E-27 | 2.645992 | 1.34E-18 |
| 13837 | 'Epha3' | 3.588964 | 1.54E-49 | 1.530071 | 1.40E-18 | 4 | 5.66E-22 | 2.125531 | 7.75E-12 |
| 20563 | 'Slit2' | 1.675565 | 1.70E-15 | 2.84549 | 6.91E-30 | 1.165586 | 4.95E-07 | 2.915608 | 7.58E-22 |
| 13841 | 'Epha7' | 2.392317 | 4.13E-09 | 2.517848 | 1.98E-09 | 2.893085 | 6.22E-12 | 2.530515 | 4.18E-10 |
