## Supplementary material for "Distinct transcriptomic profile of satellite cells contributes to preservation of neuromuscular junctions in extraocular muscles of ALS mice": Figure 7-Source Data 2

**Figure 7-Source Data 2. Axon guidance related genes (KEGG) identified in EOM SC signature genes cultured in differentiation medium.** Log2FC and FDR are shown and genes are ranked according to relative abundance (TPM, from highest to lowest) in WT EOM SCs.

| Gene ID | Gene Symbol | log2 (G93A_EOM_D / G93A_Dia_D) | FDR (G93A_EOM_D / G93A_Dia_D) | log2 (G93A _EOM_D / G93A _HL_D) | FDR (G93A _EOM_D / G93A _HL_D) | log2 (WT_EOM_D / WT_Dia_D) | FDR (WT_EOM_D / WT_Dia_D) | log2 (WT_EOM_D / WT_HL_D) | FDR (WT_EOM_D / WT_HL_D) |
| --- | --- | --- | --- | --- | --- | --- | --- | --- | --- |
| 12631 | 'Cfl1' | 0.792931 | 0 | 1.322608 | 0 | 1.456835 | 0 | 1.662317 | 0 |
| 14678 | 'Gnai2' | 0.502585 | 8.36E-96 | 0.65736 | 1.35E-148 | 0.725419 | 7.17E-198 | 0.794393 | 3.77E-229 |
| 11848 | 'Rhoa' | 0.464227 | 3.88E-92 | 0.49873 | 3.14E-94 | 0.690858 | 5.39E-160 | 0.642952 | 2.02E-141 |
| 26417 | 'Mapk3' | 0.743175 | 5.63E-42 | 1.155143 | 5.45E-85 | 0.838383 | 5.04E-46 | 0.895433 | 6.71E-51 |
| 18479 | 'Pak1' | 0.561387 | 6.71E-35 | 0.55921 | 3.28E-34 | 0.59802 | 7.55E-40 | 0.587448 | 2.67E-38 |
| 14679 | 'Gnai3' | 0.79512 | 2.01E-59 | 0.91678 | 6.16E-74 | 1.027903 | 2.00E-88 | 0.961544 | 9.79E-79 |
| 18709 | 'Pik3r2' | 0.86805 | 7.47E-63 | 0.625574 | 2.10E-35 | 0.67771 | 6.14E-42 | 0.887724 | 2.63E-65 |
| 218397 | 'Rasa1' | 0.822002 | 2.25E-78 | 0.716115 | 2.76E-60 | 1.141874 | 1.26E-151 | 1.040762 | 1.03E-128 |
| 18845 | 'Plxna2' | 0.908668 | 1.06E-181 | 0.864005 | 1.32E-163 | 0.721524 | 3.94E-106 | 0.515683 | 4.73E-58 |
| 13836 | 'Epha2' | 0.938956 | 3.02E-53 | 0.921483 | 4.82E-53 | 1.435834 | 1.34E-107 | 1.35727 | 7.20E-98 |
| 14677 | 'Gnai1' | 0.88414 | 1.61E-38 | 1.469103 | 1.58E-83 | 1.587264 | 6.76E-104 | 1.172659 | 1.70E-65 |
| 319757 | 'Smo' | 1.527605 | 8.12E-144 | 1.456509 | 5.07E-136 | 1.431951 | 9.80E-102 | 1.640004 | 3.28E-120 |
| 17295 | 'Met' | 1.58737 | 2.57E-259 | 1.634386 | 1.29E-267 | 1.129679 | 1.25E-118 | 1.047601 | 4.85E-105 |
| 20315 | 'Cxcl12' | 1.84667 | 1.09E-138 | 3.937494 | 0 | 1.762547 | 1.07E-77 | 2.380984 | 1.19E-126 |
| 107449 | 'Unc5b' | 0.588623 | 3.04E-27 | 0.720274 | 8.94E-38 | 1.198287 | 2.77E-110 | 1.559566 | 1.09E-162 |
| 20361 | 'Sema7a' | 0.412953 | 9.43E-08 | 1.174431 | 2.42E-43 | 1.09149 | 4.52E-47 | 1.227166 | 2.73E-59 |
| 16653 | 'Kras' | 0.505541 | 1.20E-18 | 0.868407 | 4.79E-46 | 0.54873 | 8.92E-22 | 0.561669 | 1.56E-22 |
| 11350 | 'Abl1' | 0.784041 | 1.31E-52 | 0.985556 | 1.58E-72 | 0.603827 | 3.81E-31 | 0.742842 | 9.37E-42 |
| 13642 | 'Efnb2' | 2.483769 | 1.52E-203 | 2.970884 | 4.85E-243 | 2.628758 | 4.39E-212 | 3.241153 | 3.48E-255 |
| 20353 | 'Sema4c' | 0.643966 | 2.62E-20 | 0.942473 | 5.19E-38 | 1.033985 | 2.96E-44 | 1.111351 | 8.77E-52 |
| 20349 | 'Sema3e' | 2.570032 | 4.22E-257 | 0.696576 | 2.92E-36 | 2.932654 | 0 | 1.948992 | 1.21E-200 |
| 65254 | 'Dpysl5' | 1.9121 | 3.67E-85 | 2.835865 | 1.46E-132 | 2.859615 | 2.76E-218 | 2.812965 | 3.21E-210 |
| 19876 | 'Robo1' | 2.921097 | 0 | 2.429244 | 0 | 2.317095 | 1.66E-248 | 2.232758 | 4.84E-234 |
| 18186 | 'Nrp1' | 2.286881 | 1.26E-177 | 2.78099 | 3.55E-225 | 2.461529 | 2.88E-183 | 2.797956 | 1.88E-210 |
| 18208 | 'Ntn1' | 1.43796 | 4.38E-112 | 1.526269 | 1.16E-119 | 1.073882 | 2.67E-53 | 1.456902 | 1.09E-84 |
| 57764 | 'Ntn4' | 1.909573 | 1.67E-91 | 2.846988 | 7.60E-144 | 0.876978 | 4.22E-16 | 1.015119 | 4.52E-20 |
| 12767 | 'Cxcr4' | 2.933289 | 1.97E-80 | 1.988518 | 4.10E-50 | 2.159446 | 1.48E-41 | 1.497354 | 2.08E-25 |
| 13846 | 'Ephb4' | 1.294542 | 4.16E-60 | 1.431553 | 2.26E-68 | 1.19977 | 1.30E-43 | 1.214125 | 4.34E-44 |
| 108151 | 'Sema3d' | 2.75086 | 2.76E-214 | 0.610539 | 1.48E-22 | 2.277305 | 3.49E-132 | 1.248305 | 8.27E-57 |
| 50780 | 'Rgs3' | 0.929527 | 2.05E-11 | 1.169925 | 3.79E-18 | 1.431105 | 4.29E-22 | 1.661162 | 6.01E-30 |
| 13835 | 'Epha1' | 2.524662 | 6.34E-75 | 3.792767 | 1.20E-107 | 3.306498 | 1.08E-94 | 4.575685 | 8.56E-120 |
| 13848 | 'Ephb6' | 4.044394 | 1.41E-103 | 2.076103 | 1.16E-51 | 3.516765 | 2.51E-105 | 3.011973 | 4.56E-92 |
| 18708 | 'Pik3r1' | 1.412843 | 3.58E-65 | 1.430491 | 2.58E-65 | 1.332196 | 1.34E-55 | 1.173767 | 5.28E-47 |
| 73181 | 'Nfatc4' | 2.886452 | 3.24E-84 | 4.954196 | 3.32E-122 | 3.208399 | 4.28E-76 | 5.025535 | 3.55E-106 |
| 13639 | 'Efna4' | 1.003336 | 2.99E-05 | 0.827486 | 5.18E-04 | 1.735353 | 1.39E-14 | 1.952382 | 8.43E-17 |
| 20352 | 'Sema4b' | 0.793667 | 6.91E-15 | 0.658683 | 8.57E-11 | 0.55011 | 3.21E-06 | 0.680507 | 9.97E-09 |
| 117606 | 'Boc' | 1.498251 | 1.01E-30 | 1.602948 | 5.34E-33 | 0.948518 | 1.89E-14 | 1.530821 | 4.23E-29 |
| 12934 | 'Dpysl2' | 1.377602 | 2.60E-30 | 2.274916 | 4.23E-58 | 1.601641 | 2.03E-31 | 3.027906 | 1.03E-66 |
| 20779 | 'Src' | 0.788496 | 4.09E-09 | 0.986007 | 5.53E-13 | 0.994051 | 2.68E-13 | 0.88544 | 6.06E-11 |
| 20346 | 'Sema3a' | 1.627523 | 8.21E-46 | 0.64054 | 1.07E-09 | 1.236819 | 4.11E-29 | 1.303589 | 2.36E-30 |
| 18481 | 'Pak3' | 1.470142 | 4.36E-51 | 2.371132 | 9.75E-94 | 1.872402 | 1.39E-67 | 1.641918 | 1.25E-54 |
| 20563 | 'Slit2' | 3.140233 | 2.24E-146 | 5.047124 | 1.36E-201 | 3.195862 | 7.86E-128 | 4.562644 | 3.57E-165 |
| 20359 | 'Sema6b' | 1.729352 | 3.66E-28 | 1.119299 | 9.47E-15 | 0.69159 | 4.93E-06 | 0.977747 | 5.34E-10 |
| 18707 | 'Pik3cd' | 1.109624 | 1.07E-11 | 1.332017 | 7.40E-15 | 0.711042 | 2.33E-05 | 0.720958 | 3.11E-05 |
| 12167 | 'Bmpr1b' | 1.82103 | 2.54E-14 | 0.72792 | 9.85E-04 | 2 | 2.14E-21 | 2.054448 | 2.29E-21 |
| 13838 | 'Epha4' | 1.90357 | 1.58E-24 | 1.222392 | 1.27E-12 | 1.135978 | 1.89E-10 | 1.054448 | 2.58E-09 |
| 20356 | 'Sema5a' | 2.61891 | 4.16E-46 | 1.068713 | 1.06E-13 | 3.841302 | 1.48E-68 | 1.966833 | 1.50E-33 |
| 22418 | 'Wnt5a' | 1.338323 | 6.26E-09 | 1.609625 | 2.63E-11 | 2.385654 | 2.71E-18 | 1.017922 | 5.45E-07 |
| 13640 | 'Efna5' | 1.534336 | 7.28E-12 | 3 | 2.79E-25 | 2.691162 | 1.31E-21 | 2.632268 | 1.04E-20 |
| 13837 | 'Epha3' | 3.394684 | 8.69E-40 | 2.895113 | 1.58E-33 | 3.447459 | 1.42E-16 | 2.736966 | 4.60E-13 |
| 22253 | 'Unc5c' | 3.820179 | 4.93E-39 | 3.235216 | 5.88E-33 | 5.196397 | 9.18E-45 | 2.974005 | 1.14E-29 |
| 268902 | 'Robo2' | 3.30117 | 2.20E-14 | 1.938599 | 2.61E-07 | 3.446256 | 2.30E-25 | 0.910203 | 1.44E-05 |
| 241568 | 'Lrrc4c' | 2.473931 | 8.37E-04 | 2.473931 | 3.36E-05 | 3.550197 | 2.22E-13 | 2.77259 | 2.24E-10 |
| 13841 | 'Epha7' | 3.169925 | 7.51E-10 | 2.491853 | 6.40E-11 | 2.920566 | 1.14E-10 | 4.72792 | 1.31E-14 |
