## Supplementary material for "Distinct transcriptomic profile of satellite cells contributes to preservation of neuromuscular junctions in extraocular muscles of ALS mice": Figure 7-Source Data 3

**Figure 7-Source Data 3. Axon guidance related genes (KEGG) identified in NaBu treatment signature genes.** Log2FC and FDR are shown and genes are ranked according to relative abundance (TPM, from highest to lowest) in G93A HL SCs with 3-day NaBu treatment.

| Gene ID | Gene Symbol | log2 (G93A_Dia_D_NaBu3 / G93A_Dia_D) | FDR (G93A_Dia_D_NaBu3 / G93A_Dia_D) | log2 (G93A_HL_D_NaBu3 / G93A_HL_D) | FDR (G93A_HL_D_NaBu3 / G93A_HL_D) |
| --- | --- | --- | --- | --- | --- |
| 12631 | 'Cfl1' | 0.465165 | 2.17E-67 | 0.536565 | 6.94E-96 |
| 13800 | 'Enah' | 0.597746 | 0 | 0.421052 | 1.86E-172 |
| 226251 | 'Ablim1' | 0.47712 | 9.73E-119 | 0.513076 | 7.42E-161 |
| 319713 | 'Ablim3' | -2.54606 | 0 | -1.33558 | 0 |
| 18844 | 'Plxna1' | 0.84319 | 0 | 0.504581 | 2.06E-119 |
| 15461 | 'Hras' | -0.98699 | 6.64E-91 | -0.99771 | 4.64E-88 |
| 18479 | 'Pak1' | 1.026498 | 3.48E-134 | 0.855623 | 5.26E-84 |
| 13836 | 'Epha2' | 2.118514 | 0 | 2.118912 | 0 |
| 20130 | 'Rras' | 0.526306 | 5.86E-09 | 0.617674 | 2.00E-11 |
| 19056 | 'Ppp3cb' | -0.51618 | 6.33E-31 | -0.56652 | 3.59E-35 |
| 14679 | 'Gnai3' | 0.660173 | 1.54E-25 | 0.575848 | 7.82E-20 |
| 259302 | 'Srgap3' | 0.82831 | 9.35E-69 | 0.607618 | 1.07E-52 |
| 108058 | 'Camk2d' | -0.55754 | 3.69E-26 | -0.57014 | 3.92E-25 |
| 18845 | 'Plxna2' | -0.91647 | 6.14E-213 | -0.78043 | 1.07E-157 |
| 13845 | 'Ephb3' | -1.34757 | 6.69E-185 | -1.14593 | 3.39E-129 |
| 20315 | 'Cxcl12' | 2.033878 | 8.03E-101 | 2.214265 | 2.93E-86 |
| 224617 | 'Tbc1d24' | 0.409224 | 1.54E-16 | 0.438996 | 1.55E-18 |
| 20361 | 'Sema7a' | 1.805184 | 1.28E-54 | 1.858241 | 3.35E-87 |
| 20349 | 'Sema3e' | -0.99709 | 1.45E-23 | -0.44457 | 2.09E-17 |
| 228026 | 'Pdk1' | 1.12519 | 5.00E-26 | 1.258585 | 6.71E-54 |
| 17295 | 'Met' | 0.635061 | 7.00E-19 | 0.815288 | 1.87E-32 |
| 231148 | 'Ablim2' | -2.28716 | 0 | -1.99926 | 9.06E-273 |
| 50780 | 'Rgs3' | 1.936435 | 1.62E-22 | 1.976054 | 1.71E-36 |
| 13637 | 'Efna2' | 0.583375 | 5.86E-06 | 0.718012 | 1.17E-07 |
| 20360 | 'Sema6c' | -2.32784 | 0 | -2.37531 | 0 |
| 57764 | 'Ntn4' | 1.079727 | 3.07E-14 | 0.528752 | 3.44E-05 |
| 19876 | 'Robo1' | 2.221817 | 3.91E-138 | 1.725967 | 6.36E-95 |
| 20347 | 'Sema3b' | -0.6258 | 3.80E-14 | -1.2256 | 2.51E-47 |
| 13846 | 'Ephb4' | 0.768517 | 1.85E-13 | 0.805857 | 2.21E-15 |
| 14270 | 'Srgap2' | -1.6194 | 7.84E-185 | -1.25963 | 1.21E-114 |
| 16885 | 'Limk1' | 0.579934 | 5.79E-07 | 0.563376 | 9.99E-06 |
| 117600 | 'Srgap1' | -2.14191 | 4.54E-149 | -1.50992 | 1.92E-159 |
| 20779 | 'Src' | 0.942389 | 1.27E-13 | 0.619472 | 3.98E-07 |
| 235611 | 'Plxnb1' | -0.87257 | 5.49E-33 | -0.81921 | 1.92E-32 |
| 12323 | 'Camk2b' | -1.40771 | 1.27E-47 | -1.37467 | 2.93E-44 |
| 14677 | 'Gnai1' | -1.14271 | 5.13E-20 | -0.80084 | 8.65E-10 |
| 20348 | 'Sema3c' | -1.29027 | 2.01E-32 | -1.29109 | 2.72E-41 |
| 26456 | 'Sema4g' | 1.5286 | 6.79E-15 | 1.79518 | 1.84E-19 |
| 73181 | 'Nfatc4' | 2.863498 | 1.77E-29 | 3.163746 | 1.07E-29 |
| 107448 | 'Unc5a' | -0.79591 | 4.82E-08 | -0.58077 | 2.23E-04 |
| 12934 | 'Dpysl2' | 1.049631 | 1.49E-07 | 1.250543 | 1.29E-09 |
| 223881 | 'Rnd1' | 2.541146 | 1.30E-13 | 2.527247 | 1.83E-11 |
| 13848 | 'Ephb6' | 3.078003 | 2.12E-09 | 1.366782 | 1.52E-08 |
| 13638 | 'Efna3' | 1.929611 | 3.67E-06 | 2.243926 | 4.29E-07 |
| 18019 | 'Nfatc2' | -0.82468 | 9.95E-17 | -0.93289 | 4.25E-10 |
| 68423 | 'Ankrd13d' | 1.642843 | 2.84E-06 | 1.417991 | 7.90E-05 |
| 65254 | 'Dpysl5' | -2.76939 | 7.40E-110 | -2.91634 | 4.43E-167 |
| 214968 | 'Sema6d' | -1.66073 | 5.26E-20 | -1.60962 | 6.03E-34 |
| 22417 | 'Wnt4' | -0.7129 | 4.52E-04 | -0.93974 | 9.95E-06 |
| 16728 | 'L1cam' | -1.77024 | 3.13E-26 | -1.88136 | 7.38E-20 |
| 214230 | 'Pak6' | 2.184425 | 2.73E-05 | 2.263034 | 2.96E-05 |
| 270190 | 'Ephb1' | -2.28758 | 1.69E-09 | -1.86876 | 1.50E-07 |
| 20562 | 'Slit1' | -1.40439 | 1.46E-04 | -1.30666 | 1.60E-05 |
| 22065 | 'Trpc3' | -2.97982 | 1.67E-09 | -3.12338 | 3.66E-07 |
| 243743 | 'Plxna4' | -2.32193 | 1.04E-19 | -2.74723 | 1.30E-17 |
