## Supplementary material for "Distinct transcriptomic profile of satellite cells contributes to preservation of neuromuscular junctions in extraocular muscles of ALS mice": Figure 7-Source Data 4

**Figure 7-Source Data 4. qRT-PCR results for *Hmga2*, *Actn3*, *Notch3* and *Cxcl12* in SCs of different muscle origins cultured in growth medium.** The averaged dCt values of the genes of interest of WT HL SCs were used as the normalization controls during the calculation of ddCt.

| Batch ID | Gender | Muscle | Genotype | Group | RQ_*Actn3* | RQ_*Cxcl12* | RQ_*Hmga2* | RQ_*Notch3* |
| --- | --- | --- | --- | --- | --- | --- | --- | --- |
| WT-HL-1 | M | HL | WT | Growth | 2.190774 | 0.965134 | 0.515255 | 0.471589 |
| WT-HL-1 | M | HL | WT | Growth | 2.227721 | 0.976944 | 0.536088 | 0.558002 |
| WT-HL-1 | M | HL | WT | Growth | 2.269848 | 0.925279 | 0.467634 | 0.51919 |
| WT-HL-2 | M | HL | WT | Growth | 1.551981 | 0.426435 | 0.796629 | 0.611102 |
| WT-HL-2 | M | HL | WT | Growth | 1.437594 | 0.431597 | 0.794492 | 0.605704 |
| WT-HL-2 | M | HL | WT | Growth | 1.62728 | 0.425993 | 0.843011 | 0.655562 |
| WT-HL-3 | M | HL | WT | Growth | 0.202171 | 4.062814 | 1.912532 | 0.973669 |
| WT-HL-3 | M | HL | WT | Growth | 0.203709 | 4.213902 | 2.165718 | 1.029941 |
| WT-HL-3 | M | HL | WT | Growth | 0.195057 | 4.205592 | 2.047402 | 1.300646 |
| WT-HL-4 | F | HL | WT | Growth | 1.39026 | 0.552182 | 1.262922 | 2.767921 |
| WT-HL-4 | F | HL | WT | Growth | 1.449346 | 0.605161 | 1.191514 | 2.993201 |
| WT-HL-4 | F | HL | WT | Growth | 1.536035 | 0.607636 | 1.13702 | 2.791345 |
| WT_Dia-1 | F | Dia | WT | Growth | 4.502679 | 2.333534 | 0.555072 | 1.845976 |
| WT_Dia-1 | F | Dia | WT | Growth | 4.06114 | 2.06765 | 0.543321 | 2.066691 |
| WT_Dia-1 | F | Dia | WT | Growth | 3.856546 | 1.892705 | 0.416696 | 2.761623 |
| WT_Dia-2 | M | Dia | WT | Growth | 1.235518 | 3.043082 | 0.587296 | 0.970943 |
| WT_Dia-2 | M | Dia | WT | Growth | 1.19574 | 2.923508 | 0.528997 | 0.913106 |
| WT_Dia-2 | M | Dia | WT | Growth | 1.246233 | 2.756881 | 0.533323 | 0.888613 |
| WT_Dia-3 | M | Dia | WT | Growth | 3.543202 | 1.830473 | 0.889786 | 0.591017 |
| WT_Dia-3 | M | Dia | WT | Growth | 3.280003 | 1.817959 | 0.847683 | 0.514347 |
| WT_Dia-3 | M | Dia | WT | Growth | 3.383235 | 1.780719 | 0.85137 | 0.524983 |
| WT_Dia-4 | F | Dia | WT | Growth | 5.107195 | 3.237634 | 1.352536 | 2.292361 |
| WT_Dia-4 | F | Dia | WT | Growth | 4.874007 | 2.927712 | 1.306793 | 2.093935 |
| WT_Dia-4 | F | Dia | WT | Growth | 5.072299 | 2.992849 | 1.322548 | 2.149072 |
| WT-EOM-1 | M | EOM | WT | Growth | 0.306175 | 7.986597 | 5.058652 | 0.378471 |
| WT-EOM-1 | M | EOM | WT | Growth | 0.280947 | 7.843129 | 5.313471 | 0.479953 |
| WT-EOM-1 | M | EOM | WT | Growth | 0.259006 | 7.684834 | 3.702898 | 0.414803 |
| WT-EOM-2 | M | EOM | WT | Growth | 1.423388 | 7.367453 | 2.028652 | 0.997135 |
| WT-EOM-2 | M | EOM | WT | Growth | 1.388814 | 7.108022 | 1.860727 | 0.915022 |
| WT-EOM-2 | M | EOM | WT | Growth | 1.382522 | 7.398815 | 1.872449 | 0.957845 |
| WT-EOM-3 | M | EOM | WT | Growth | 1.297522 | 7.711204 | 2.375671 | 3.387115 |
| WT-EOM-3 | M | EOM | WT | Growth | 1.280303 | 7.477761 | 2.123877 | 2.895819 |
| WT-EOM-3 | M | EOM | WT | Growth | 1.142566 | 7.667197 | 2.086641 | 4.783497 |
| G93A-HL-1 | F | HL | G93A | Growth | 0.613727 | 1.500434 | 1.29308 | 0.60758 |
| G93A-HL-1 | F | HL | G93A | Growth | 0.592415 | 1.704578 | 1.448764 | 0.659259 |
| G93A-HL-1 | F | HL | G93A | Growth | 0.588781 | 1.575128 | 1.298325 | 0.610556 |
| G93A-HL-2 | M | HL | G93A | Growth | 3.836236 | 1.640624 | 1.15731 | 1.493616 |
| G93A-HL-2 | M | HL | G93A | Growth | 4.143258 | 1.704713 | 1.178559 | 1.543202 |
| G93A-HL-2 | M | HL | G93A | Growth | 2.965006 | 1.71149 | 1.196098 | 1.541283 |
| G93A-HL-3 | M | HL | G93A | Growth | 0.162281 | 7.290182 | 2.585049 | 0.315422 |
| G93A-HL-3 | M | HL | G93A | Growth | 0.194567 | 6.800504 | 2.147042 | 0.317433 |
| G93A-HL-3 | M | HL | G93A | Growth | 0.197992 | 7.585659 | 2.371438 | 0.409719 |
| G93A-HL-4 | M | HL | G93A | Growth | 1.754546 | 2.65703 | 0.417143 | 1.290678 |
| G93A-HL-4 | M | HL | G93A | Growth | 2.012665 | 3.024939 | 0.493979 | 1.545652 |
| G93A-HL-4 | M | HL | G93A | Growth | 1.713788 | 2.306085 | 0.389875 | 2.099488 |
| G93A-HL-5 | M | HL | G93A | Growth | 0.31073 | 9.541869 | 1.275026 | 0.703097 |
| G93A-HL-5 | M | HL | G93A | Growth | 0.306103 | 9.614174 | 1.239493 | 0.703839 |
| G93A-HL-5 | M | HL | G93A | Growth | 0.316929 | 9.056518 | 1.157429 | 0.72583 |
| G93A-HL-6 | F | HL | G93A | Growth | 0.239735 | 3.894275 | 1.645712 | 1.056748 |
| G93A-HL-6 | F | HL | G93A | Growth | 0.266863 | 3.99096 | 1.655933 | 1.083439 |
| G93A-HL-6 | F | HL | G93A | Growth | 0.289601 | 4.069674 | 1.752435 | 1.064961 |
| G93A-Dia-1 | F | Dia | G93A | Growth | 0.510699 | 6.945521 | 0.607705 | 0.891473 |
| G93A-Dia-1 | F | Dia | G93A | Growth | 0.549587 | 7.015873 | 0.655753 | 0.984867 |
| G93A-Dia-1 | F | Dia | G93A | Growth | 0.487478 | 7.312306 | 0.795546 | 1.026554 |
| G93A-Dia-2 | F | Dia | G93A | Growth | 1.4637 | 10.37355 | 2.388852 | 0.752258 |
| G93A-Dia-2 | F | Dia | G93A | Growth | 1.470741 | 10.48042 | 2.405334 | 0.733088 |
| G93A-Dia-2 | F | Dia | G93A | Growth | 1.442863 | 10.14226 | 2.401283 | 0.69574 |
| G93A-Dia-3 | F | Dia | G93A | Growth | 1.284278 | 3.227792 | 1.088593 | 0.574383 |
| G93A-Dia-3 | F | Dia | G93A | Growth | 1.242981 | 3.102838 | 1.06766 | 0.520693 |
| G93A-Dia-3 | F | Dia | G93A | Growth | 1.300135 | 3.042957 | 1.036396 | 0.515049 |
| G93A-Dia-4 | M | Dia | G93A | Growth | 1.435796 | 0.685468 | 0.472373 | 0.74247 |
| G93A-Dia-4 | M | Dia | G93A | Growth | 1.448543 | 0.667923 | 0.515516 | 0.724029 |
| G93A-Dia-4 | M | Dia | G93A | Growth | 1.427543 | 0.733253 | 0.480565 | 0.657373 |
| G93A-EOM-1 | M | EOM | G93A | Growth | 0.973947 | 7.923536 | 3.669496 | 0.637045 |
| G93A-EOM-1 | M | EOM | G93A | Growth | 0.949916 | 8.274177 | 3.590819 | 0.630245 |
| G93A-EOM-1 | M | EOM | G93A | Growth | 0.883771 | 8.198542 | 3.313482 | 0.613837 |
| G93A-EOM-2 | M | EOM | G93A | Growth | 0.102719 | 5.026044 | 1.68864 | 0.203328 |
| G93A-EOM-2 | M | EOM | G93A | Growth | 0.098408 | 5.24308 | 1.62561 | 0.203304 |
| G93A-EOM-2 | M | EOM | G93A | Growth | 0.116023 | 5.218206 | 1.641714 | 0.254192 |
| G93A-EOM-3 | M | EOM | G93A | Growth | 0.094592 | 11.79538 | 2.288458 | 0.833387 |
| G93A-EOM-3 | M | EOM | G93A | Growth | 0.095306 | 12.17947 | 2.239922 | 0.782778 |
| G93A-EOM-3 | M | EOM | G93A | Growth | 0.095411 | 12.21986 | 2.120023 | 0.814051 |
