## Supplementary material for "Distinct transcriptomic profile of satellite cells contributes to preservation of neuromuscular junctions in extraocular muscles of ALS mice": Figure 7-Source Data 6

**Figure 7-Source Data 6. qRT-PCR results for *Hmga2*, *Actn3*, *Notch3* and *Cxcl12* in G93A diaphragm and HL SCs with or without 3-day NaBu treatment.** The averaged dCt values of the genes of interest of G93A HL SCs were used as the normalization controls during the calculation of ddCt.

| Batch ID | Gender | Muscle | Genotype | Group | RQ_*Actn3* | RQ_*Cxcl12* | RQ_*Hmga2* | RQ_*Notch3* |
| --- | --- | --- | --- | --- | --- | --- | --- | --- |
| G93A-Dia-1 | F | Dia | G93A | Differentiation | 0.87176 | 2.734384 | 1.081501 | 1.401929 |
| G93A-Dia-1 | F | Dia | G93A | Differentiation | 0.87435 | 2.596057 | 0.986736 | 2.061618 |
| G93A-Dia-1 | F | Dia | G93A | Differentiation | 0.958265 | 3.176493 | 1.011654 | 1.831684 |
| G93A-Dia-1 | F | Dia | G93A | Differentiation_NaBu3 | 0.517017 | 10.84023 | 2.358103 | 4.811868 |
| G93A-Dia-1 | F | Dia | G93A | Differentiation_NaBu3 | 0.570675 | 11.30371 | 2.454603 | 4.358444 |
| G93A-Dia-1 | F | Dia | G93A | Differentiation_NaBu3 | 0.536912 | 11.07001 | 2.095331 | 4.289764 |
| G93A-Dia-2 | F | Dia | G93A | Differentiation | 1.401197 | 0.830592 | 1.664961 | 0.953365 |
| G93A-Dia-2 | F | Dia | G93A | Differentiation | 1.355486 | 0.716768 | 1.527939 | 0.98458 |
| G93A-Dia-2 | F | Dia | G93A | Differentiation | 1.450161 | 0.90436 | 1.688148 | 0.966295 |
| G93A-Dia-2 | F | Dia | G93A | Differentiation_NaBu3 | 0.877314 | 2.761066 | 0.850886 | 3.952817 |
| G93A-Dia-2 | F | Dia | G93A | Differentiation_NaBu3 | 0.861309 | 2.649013 | 0.870316 | 3.904993 |
| G93A-Dia-2 | F | Dia | G93A | Differentiation_NaBu3 | 0.827272 | 2.655373 | 0.850391 | 3.953203 |
| G93A-Dia-3 | F | Dia | G93A | Differentiation | 0.833287 | 0.741925 | 0.678712 | 0.434045 |
| G93A-Dia-3 | F | Dia | G93A | Differentiation | 0.79259 | 0.846085 | 0.643368 | 0.468727 |
| G93A-Dia-3 | F | Dia | G93A | Differentiation | 0.788755 | 0.85568 | 0.709863 | 0.45607 |
| G93A-Dia-3 | F | Dia | G93A | Differentiation_NaBu3 | 0.486465 | 5.287879 | 3.090082 | 3.187678 |
| G93A-Dia-3 | F | Dia | G93A | Differentiation_NaBu3 | 0.490959 | 5.564999 | 3.078664 | 3.044019 |
| G93A-Dia-3 | F | Dia | G93A | Differentiation_NaBu3 | 0.456051 | 5.300288 | 3.044978 | 3.000722 |
| G93A-Dia-4 | M | Dia | G93A | Differentiation | 0.811886 | 0.270001 | 0.855366 | 0.474089 |
| G93A-Dia-4 | M | Dia | G93A | Differentiation | 0.711539 | 0.268824 | 0.859294 | 0.510407 |
| G93A-Dia-4 | M | Dia | G93A | Differentiation | 0.722399 | 0.277502 | 0.930112 | 0.503623 |
| G93A-Dia-4 | M | Dia | G93A | Differentiation_NaBu3 | 0.622415 | 0.421276 | 2.756991 | 4.053388 |
| G93A-Dia-4 | M | Dia | G93A | Differentiation_NaBu3 | 0.600826 | 0.392929 | 2.674161 | 3.794911 |
| G93A-Dia-4 | M | Dia | G93A | Differentiation_NaBu3 | 0.614088 | 0.447593 | 2.707526 | 3.677848 |
| G93A-Dia-5 | M | Dia | G93A | Differentiation | 1.073232 | 1.3813 | 0.664055 | 1.812295 |
| G93A-Dia-5 | M | Dia | G93A | Differentiation | 1.113936 | 1.428132 | 0.711327 | 1.509928 |
| G93A-Dia-5 | M | Dia | G93A | Differentiation | 1.141179 | 1.445823 | 0.615922 | 1.718217 |
| G93A-Dia-5 | M | Dia | G93A | Differentiation_NaBu3 | 0.709995 | 6.163537 | 1.737568 | 6.921953 |
| G93A-Dia-5 | M | Dia | G93A | Differentiation_NaBu3 | 0.770444 | 8.107937 | 1.781225 | 6.484392 |
| G93A-Dia-5 | M | Dia | G93A | Differentiation_NaBu3 | 0.782367 | 7.723788 | 1.773073 | 7.206772 |
| G93A-Dia-6 | F | Dia | G93A | Differentiation | 1.13831 | 1.305539 | 1.518675 | 1.598659 |
| G93A-Dia-6 | F | Dia | G93A | Differentiation | 1.143011 | 1.272529 | 1.576103 | 1.303572 |
| G93A-Dia-6 | F | Dia | G93A | Differentiation | 1.288089 | 1.60682 | 1.461588 | 1.879621 |
| G93A-Dia-6 | F | Dia | G93A | Differentiation_NaBu3 | 0.824569 | 2.633272 | 1.233367 | 3.781284 |
| G93A-Dia-6 | F | Dia | G93A | Differentiation_NaBu3 | 0.787219 | 2.423135 | 1.16116 | 3.693558 |
| G93A-Dia-6 | F | Dia | G93A | Differentiation_NaBu3 | 0.987033 | 2.77807 | 1.28614 | 4.045812 |
| G93A-HL-1 | M | HL | G93A | Differentiation | 0.864598 | 1.527541 | 1.759258 | 1.47022 |
| G93A-HL-1 | M | HL | G93A | Differentiation | 0.889222 | 1.522021 | 1.719259 | 1.342572 |
| G93A-HL-1 | M | HL | G93A | Differentiation | 0.962773 | 1.369403 | 1.338486 | 1.494758 |
| G93A-HL-1 | M | HL | G93A | Differentiation_NaBu3 | 0.470092 | 21.89259 | 16.00955 | 5.791899 |
| G93A-HL-1 | M | HL | G93A | Differentiation_NaBu3 | 0.497014 | 23.97474 | 13.71168 | 5.951595 |
| G93A-HL-1 | M | HL | G93A | Differentiation_NaBu3 | 0.449822 | 21.14783 | 12.13631 | 6.473635 |
| G93A-HL-2 | M | HL | G93A | Differentiation | 1.295463 | 0.581481 | 1.070545 | 0.469536 |
| G93A-HL-2 | M | HL | G93A | Differentiation | 1.387882 | 0.57184 | 1.009161 | 0.504434 |
| G93A-HL-2 | M | HL | G93A | Differentiation | 1.339882 | 0.601182 | 0.895202 | 0.464335 |
| G93A-HL-2 | M | HL | G93A | Differentiation_NaBu3 | 0.712388 | 4.036013 | 3.798408 | 2.436057 |
| G93A-HL-2 | M | HL | G93A | Differentiation_NaBu3 | 0.909558 | 3.916553 | 3.725084 | 2.14895 |
| G93A-HL-2 | M | HL | G93A | Differentiation_NaBu3 | 0.929896 | 4.0428 | 3.670667 | 2.306022 |
| G93A-HL-3 | M | HL | G93A | Differentiation | 0.752227 | 0.725583 | 0.56245 | 0.99502 |
| G93A-HL-3 | M | HL | G93A | Differentiation | 0.736138 | 0.75816 | 0.564186 | 0.950717 |
| G93A-HL-3 | M | HL | G93A | Differentiation | 0.729426 | 0.703196 | 0.521057 | 1.049677 |
| G93A-HL-3 | M | HL | G93A | Differentiation_NaBu3 | 0.533607 | 4.049161 | 1.632048 | 2.93527 |
| G93A-HL-3 | M | HL | G93A | Differentiation_NaBu3 | 0.506559 | 3.775633 | 1.449064 | 1.429638 |
| G93A-HL-3 | M | HL | G93A | Differentiation_NaBu3 | 0.527083 | 3.994374 | 1.518303 | 6.197757 |
| G93A-HL-4 | F | HL | G93A | Differentiation | 0.874134 | 1.313026 | 0.788117 | 0.931534 |
| G93A-HL-4 | F | HL | G93A | Differentiation | 0.950028 | 1.198321 | 0.771465 | 0.962419 |
| G93A-HL-4 | F | HL | G93A | Differentiation | 0.997552 | 1.223342 | 0.792585 | 1.565647 |
| G93A-HL-4 | F | HL | G93A | Differentiation_NaBu3 | 0.956679 | 6.664818 | 1.318436 | 8.46733 |
| G93A-HL-4 | F | HL | G93A | Differentiation_NaBu3 | 0.967376 | 5.810579 | 1.400668 | 8.049311 |
| G93A-HL-4 | F | HL | G93A | Differentiation_NaBu3 | 1.096667 | 6.350738 | 1.444475 | 15.25247 |
| G93A-HL-5 | M | HL | G93A | Differentiation | 1.087753 | 1.717537 | 2.702218 | 1.756909 |
| G93A-HL-5 | M | HL | G93A | Differentiation | 1.089333 | 2.965126 | 2.09319 | 1.493251 |
| G93A-HL-5 | M | HL | G93A | Differentiation | 1.009683 | 2.390603 | 2.371859 | 1.738869 |
| G93A-HL-5 | M | HL | G93A | Differentiation_NaBu3 | 0.692305 | 19.57262 | 13.25071 | 15.74086 |
| G93A-HL-5 | M | HL | G93A | Differentiation_NaBu3 | 0.651309 | 22.93493 | 10.36822 | 12.04181 |
| G93A-HL-5 | M | HL | G93A | Differentiation_NaBu3 | 0.873216 | 23.17971 | 10.90866 | 12.56822 |
| G93A-HL-6 | F | HL | G93A | Differentiation | 1.131782 | 0.620642 | 0.64678 | 0.74356 |
| G93A-HL-6 | F | HL | G93A | Differentiation | 1.125445 | 0.534014 | 0.60734 | 0.837126 |
| G93A-HL-6 | F | HL | G93A | Differentiation | 1.099775 | 0.522961 | 0.608243 | 0.778672 |
| G93A-HL-6 | F | HL | G93A | Differentiation_NaBu3 | 0.735451 | 2.283687 | 1.57682 | 2.166731 |
| G93A-HL-6 | F | HL | G93A | Differentiation_NaBu3 | 0.773955 | 2.236384 | 1.775805 | 2.61193 |
| G93A-HL-6 | F | HL | G93A | Differentiation_NaBu3 | 0.75375 | 2.213087 | 1.617982 | 3.766616 |
